# A proteomic map of B cell activation and its shaping by mTORC1, MYC and iron

**DOI:** 10.1101/2024.12.19.629506

**Authors:** Olivia James, Linda V. Sinclair, Nadejda Lefter, Anna A.C. Brock, Fiamma Salerno, Alejandro Brenes, Hanif J. Khameneh, Matteo Pecoraro, Greta Guarda, Andrew J.M. Howden

## Abstract

Using high resolution quantitative mass spectrometry, we have explored how immune activation and the metabolic checkpoint kinase mTORC1 (mammalian target of rapamycin complex 1) regulate the proteome of B lymphocytes. B cell activation via the B cell receptor, CD40 and the IL4 receptor induced considerable re-modelling of the B cell protein landscape, with a 5-fold increase in total cellular protein mass within 24 hours of activation. Analysis of copy numbers per cell of >7,500 proteins revealed increases in the metabolic machinery that supports B cell activation and nutrient and amino acid transporters that fuel B cell biosynthetic capacity. Comparison of T cell-dependent versus T cell-independent B cell stimulation along with an analysis of the synergistic effects of stimuli, identified shared and unique impacts of stimuli and highlighted the drivers of B cell proteome remodelling. In addition, we reveal that mTORC1 controls activation-induced B cell growth and inhibiting mTORC1 impairs the expression of amino acid transporters that fuel B cell protein production. We also show that mTORC1 activity regulates the expression of the transcription factor MYC and the transferrin receptor CD71. Blocking MYC activity phenocopied mTORC1 inhibition in many ways including impaired CD71 expression, while limiting iron availability during B cell activation impaired B cell growth and protein synthesis. This work provides a detailed map of naïve and immune activated B cell proteomes and a greater understanding of the cellular machinery that directs B cell phenotypes. This work also provides new insights into the role of mTORC1, MYC and iron in regulating activation-induced proteome remodelling and protein production in B cells.

## Introduction

B lymphocytes (B cells) are a key component of the adaptive immune system, producing antibodies for targeting bacteria and viruses, secreting pro- and anti-inflammatory cytokines and presenting antigen to T cells. B cell function is critical for maintaining health and as such B cell dysfunction is linked to a broad spectrum of diseases including autoimmunity and immunodeficiency, diseases of the central nervous system and B cell malignancies. B cells sense and respond to their environment using the B cell receptor (BCR) and a range of other cell surface receptors including CD40, which binds to CD40 ligand (CD40L) expressed on the surface of T cells, the pattern recognition Toll-Like Receptors including TLR4, which is triggered by lipopolysaccharide, and cytokine receptors including interleukin 4 receptor (IL4R) and interleukin 21 receptor (IL21R). Immune activation of B cells triggers transcriptional re-programming driving proliferative expansion and B cell differentiation. Effective B cell activation will give rise to populations of short and long-lived antibody producing plasma cells as well as memory B cells. Thus, activated B cells need to ramp up metabolism and protein production appropriately to sustain these diverse fates.

A central metabolic regulator that controls protein synthesis in mammalian cells is the nutrient sensing serine/threonine protein kinase complex mTORC1 (Liu & Sabatini 2020). Various studies have outlined the importance of mTORC1 in B cell growth and differentiation, placing the nutrient-regulated kinase as a critical regulator of B cell function. mTORC1 is important in B cell class switching, somatic hypermutation, memory B cell formation and the unfolded protein response (Gaudette et al. 2020a; Iwata et al. 2016, 2017; Raybuck et al. 2018; Zhang et al. 2011). However, the mechanistic details of how mTORC1 regulates B cell activation, metabolism and protein synthesis are still to be fully mapped. Given the cell-type and cell-context specific impacts of mTORC1, a detailed analysis of the role of mTORC1 in B cell activation is needed if we are to fully understand its importance during immune challenge.

Quantitative proteomics is a valuable tool to understand immune cell phenotypes and cellular responses to stimulation, stress and the modulation of critical signalling pathways(Hukelmann et al. 2016; Rieckmann et al. 2017; Salerno et al. 2023; Tan et al. 2017) . A deep characterisation of the protein landscape of the cell may help to bridge the gap between transcriptional changes and metabolic readouts, providing quantitative data on the metabolic machinery and regulators that drive cell phenotypes and function. In this study we use high sensitivity quantitative mass spectrometry to map the activated B cell proteome. We identify and quantify over 7500 proteins in naïve and activated B cells and provide estimates of absolute copy numbers per cell for each protein. We find that the naïve B cell increases its protein mass 5-fold within 24 hours of activation leading to over 5000 proteins increasing in abundance and a handful of proteins dropping in abundance including regulators of translation and cell cycle. We highlight the core processes and metabolic machinery that are regulated by B cell activation and demonstrate the critical role of the System L large neutral amino acid transporter, SLC7A5, in B cell activation. A comparative analysis of T cell-dependent versus T cell-independent stimulation on B cell proteome remodelling identifies differences in the scale of proteome remodelling along with selective impacts on protein expression. We also dissect the synergistic effect of B cell stimuli and provide new mechanistic insight into the additive impacts of B cell stimuli on metabolic machinery and proteome remodelling. To explore the metabolic regulation of B cell activation in more detail we examined the role of mTORC1, MYC and iron. mTORC1 inhibition dramatically impacts activation-induced proteome remodelling and impairs B cell protein production. mTORC1 activity is required for the sustained expression of the transcription factor MYC and for the upregulation of amino acid transporters that fuel B cell protein production, including SLC7A5. mTORC1 also regulates the expression of the transferrin receptor CD71, which is critical for iron uptake in cells. Inhibiting MYC phenocopies many of the effects of mTORC1 inhibition including the reduced expression of CD71, while limiting iron availability during B cell activation impairs B cell growth and protein synthesis. We provide new insight into how B cells respond to stimulation and modulate their core metabolic processes and machinery for fuelling protein synthesis, shedding new light on the role of mTORC1, MYC and iron during B cell activation. Our proteomics data is freely available and easy to explore on the Immunological Proteome Resource(Brenes et al. 2023) www.immpres.co.uk.

## Results

### Activation-induced proteome remodelling in B cells

To assess the intrinsic changes driven by B cell activation, high resolution quantitative mass spectrometry was used to characterise the proteomes of primary naïve and immune-activated B cells, identifying and quantifying >7500 proteins. 24hr stimulation of B cells with anti-IgM, anti-CD40 and IL4 resulted in a considerable increase in B cell size as shown by flow cytometry (Fig 1a) and a 5-fold increase in total cellular protein mass (Fig 1b). Using the proteomic ruler (Wiśniewski et al. 2014) we estimated the copy numbers per cell of all proteins identified in naïve and activated B cells. Proteins were ranked according to their estimated copy number per cell and plotted against their cumulative contribution to total protein abundance, to evaluate which proteins dominate the protein mass of naïve and activated B cells (Fig 1c). In naïve B cells, the 7 most abundant proteins which contribute 50% of the total protein mass of the cell are largely histones and cytoskeleton molecules while in activated cells 147 proteins contribute 50% of the total protein mass and includes ribosomal proteins and translational machinery, revealing that immune activation remodels the core proteome of B cells to drive protein synthesis (Fig 1c). To explore this further we analysed the contribution of different cellular compartments to the overall protein mass of the cell. Ribosomal proteins made up ∼3% of the total protein mass of naïve cells which increased to 9% following activation, while mitochondrial proteins rose from 11% to 16% (Fig 1d). Some components such as the nuclear envelope and plasma membrane, were present in higher copies following activation but when adjusted against the total cellular protein content, showed little change relative to the total proteome (Fig 1d).

**Figure 1.**
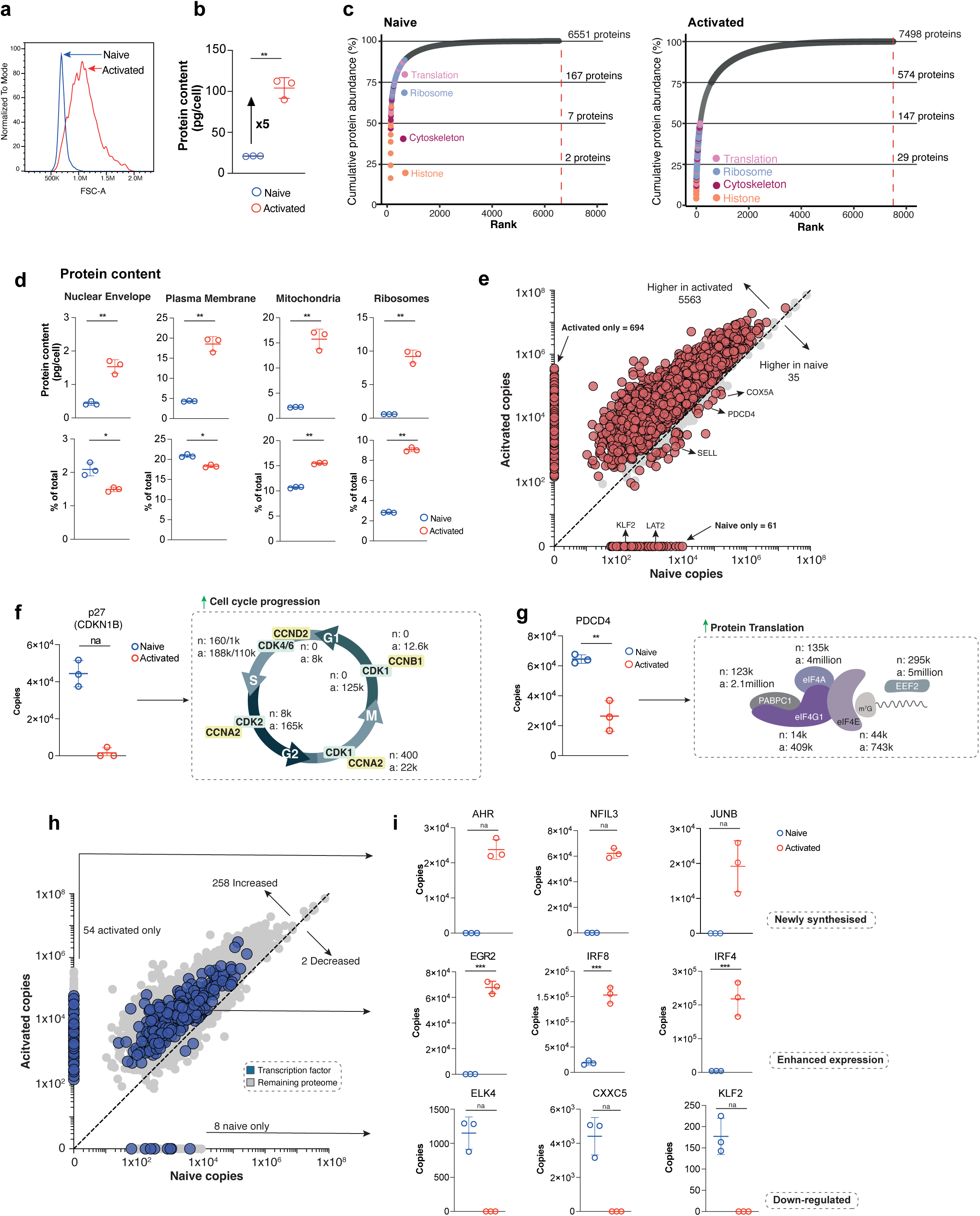
Activation-induced proteome remodelling in B cells. **(a)** Flow cytometry forward scatter (FSC) profiles of naïve B cells (blue) and 24h anti-IgM, anti-CD40 and IL4-activated B cells (red, referred to as ‘activated’). Data is representative of 3 biological replicates. **(b)** Total protein content (pg/cell) of naïve and 24h activated B cells. **(c)** Proteins ranked according to descending abundance and plotted against their cumulative abundance (shown as a percentage), for both naïve and 24h activated B cells. The number of proteins that make up each quartile of the proteome summed with the quartile below are shown on the designated section on each graph. Proteins making up the top-most 50% of activated B cells were annotated as either histones (orange), cytoskeletal (magenta), ribosomal (blue) or translational (pink) and were also mapped over the naïve B cell plot. **(d)** Top panel shows the total protein mass (pg/cell) of proteins belonging to cellular compartments including nuclear envelope (GOCC:0005635), plasma membrane (GOCC:0005886), mitochondria (GOCC:0005739) and ribosomes (KEGG 03010) as per Gene Ontology/KEGG Orthology terms. Lower panels show the % contribution to the total protein content for each term. **(e)** Differential expression analysis between naïve and activated B cells. Proteins were categorised as significantly changing between naïve and 24h activated B cells with a *P* < 0.05; fold change > 1.5 or found in one population but not detected in another population. Scatter plot shows the average protein copy number of each protein identified between naïve and 24h activated B cells. Proteins highlighted in red were significantly upregulated >1.5-fold between naïve and activated B cells, while proteins shown in blue were significantly downregulated. Proteins on the axes were found in one condition but not the other, numbers are provided on the plot for proteins found in 3/3 replicates of one condition. **(f)** Expression data for the cyclin dependent kinase inhibitor p27 (CDKN1B) and key regulators of the cell cycle. **(g)** Expression data for components of the eiF4F translational complex including the translational repressor PDCD4. **(h)**. Over 400 transcription factor and DNA-binding proteins were identified using the ‘Human transcription factor’ list (ref: https://doi.org/10.1016/j.cell.2018.01.029) as a reference database. Scatter plot compares copy number expression profiles of the transcription factors (blue) between naïve and 24h activated B cells against the total proteome (grey). Proteins were categorised as significantly changing between naïve and 24h activated B cells with a *P* < 0.05; fold change > 1.5 or found in one population but not detected in another population. Scatter plot shows the average protein copy number of each protein identified between naïve and 24h activated B cells. **(i)** Histograms showing protein copy numbers per cell for a selection of transcription factors such as those only expressed by activated B cells (newly synthesised), those with higher expression in activated as compared to naïve B cells (enhanced expression) and those only expressed by naïve B cells. For e, f, g, h and i statistical significance was derived from two-tailed empirical Bayes moderated t-statistics performed in limma on the total dataset where ** p<0.01, *** p<0.001. For b and d statistical significance was derived from an unpaired two-tailed t test where * p<0.05, ** p<0.01. All data are 3 biologically independent samples. All error bars are mean with s.d.

To examine protein changes in more detail, differential expression analysis was performed and revealed that >5000 proteins were significantly increased in expression >1.5-fold and with a p value <0.05 in activated B cells as compared to naïve (Fig 1e). The full data set along with fold changes and p values can be found in Supplementary File 1. Among those proteins increasing in abundance were activation receptors CD86, CD69 and CD44 (Supplementary File 1). 61 proteins were found exclusively in naïve B cells and not in activated B cells, suggesting that they may be degraded upon B cell stimulation. These proteins included molecules controlling B cell quiescence such as the transcription factor KLF2 (Hart et al. 2011; Winkelmann et al. 2014; Wittner & Schuh 2023) (Fig 1e). Only 35 proteins were significantly downregulated upon B cell activation. Among these downregulated proteins were negative regulators of key processes facilitating protein synthesis and proliferation such as the mRNA translational repressor PDCD4 (Loh et al. 2009; Suzuki et al. 2008) and the cell cycle inhibitor p27 (CDKN1B) (Fig 1e-g). The loss of p27 was accompanied by an increase in positive regulators of cell cycle such as cyclin-dependent kinases (CDK1-4), as well as A-type, B-type, and D-type cyclins (Fig 1f). Subunits of the EIF4F translational complex responsible for mRNA translation also showed an increase in abundance and corresponding the loss of PDCD4 (Fig 1g).

There was also considerable modulation of transcription factor expression upon B cell activation. Over 400 proteins with predicted transcription factor activity were identified within the data set (Fig 1h Supplementary File 1). Most of these transcription factors increased in abundance in response to immune activation and some were only detected in activated cells including NFIL3, AHR and JUNB (Fig 1h and i). NFIL3 expression is regulated by IL4 and is critical for B cell class switching (Kashiwada et al. 2010) . Other transcription factors were considerably upregulated upon activation, including IRF4 which increased from around 4,000 copies per cell in naïve B cells to over 200,000 copies per cell upon activation and IRF8 which increased from approximately 20,000 copies to 150,000 copies. EGR2, which controls antigen receptor induced proliferation in B cells (Li et al. 2012) , increased from around 200 copies in naïve cells to almost 70,000 copies upon activation. Some transcription factors increased by a much smaller scale including PAX5 (Supplementary File 1), which increased less than 2-fold and which has previously been shown to be important during B cell development (Cobaleda et al. 2007; Fedl et al. 2024) . A small number of transcription factors were downregulated upon activation or were exclusively found in naïve cells including ELK4, CXXC5 and KLF2 (Fig 1h and i).

### Immune activation drives nutrient transporter expression and metabolic and mitochondrial remodelling

24-hour B cell activation was accompanied by a considerable increase in cell size and total protein content. This rapid increase in cell size is energetically demanding and would require increased uptake of nutrients to fuel anabolic processes. We next asked which nutrient transporters were expressed in response to B cell activation. Several solute carrier (SLC) molecules were upregulated upon activation including amino acid transporters SLC1A5, a glutamine transporter, and SLC7A5 which transports many essential amino acids including methionine, tryptophan and leucine (Kanai et al. 1998; Scalise et al. 2018) (Fig 2a). Expression of these transporters was extremely low in naïve B cells (Fig 2a). SLC7A5 is critical for antigen-driven T cell responses, T cell differentiation and effector function (Sinclair et al. 2013, 2019) .To explore the importance of SLC7A5 in B cell activation we used splenocytes from VavCre x *Slc7a5^fl/fl^*mice (Sinclair et al. 2013) . In these mice, VavCre drives deletion of *Slc7a5* in early haematopoietic development. Splenocytes from VavCre x *Slc7a5^fl/fl^* mice were activated using anti-IgM and anti-CD40. The SLC7A5 KO B cells were considerably smaller than cre negative controls after activation (Fig 2b). SLC7A5 KO B cells also showed reduced expression of CD98/SLC3A2, the heavy chain chaperone which forms a heterodimer with many amino acid transporters, including SLC7A5 and SLC7A6 (Palacín & Kanai 2004) . Additionally, SLC7A5 KO B cells had impaired expression of the transferrin receptor CD71 in response to immune activation. CD71 is essential for iron uptake which fuels multiple metabolic processes in B cells (Frost et al. 2020; Jabara et al. 2016; Neckers et al. 1984) (Fig 2b). Together this data highlights the important role of a single amino acid transporter in fuelling the increase in size and the upregulation of key proteins that fuel B cell immune activation.

**Figure 2.**
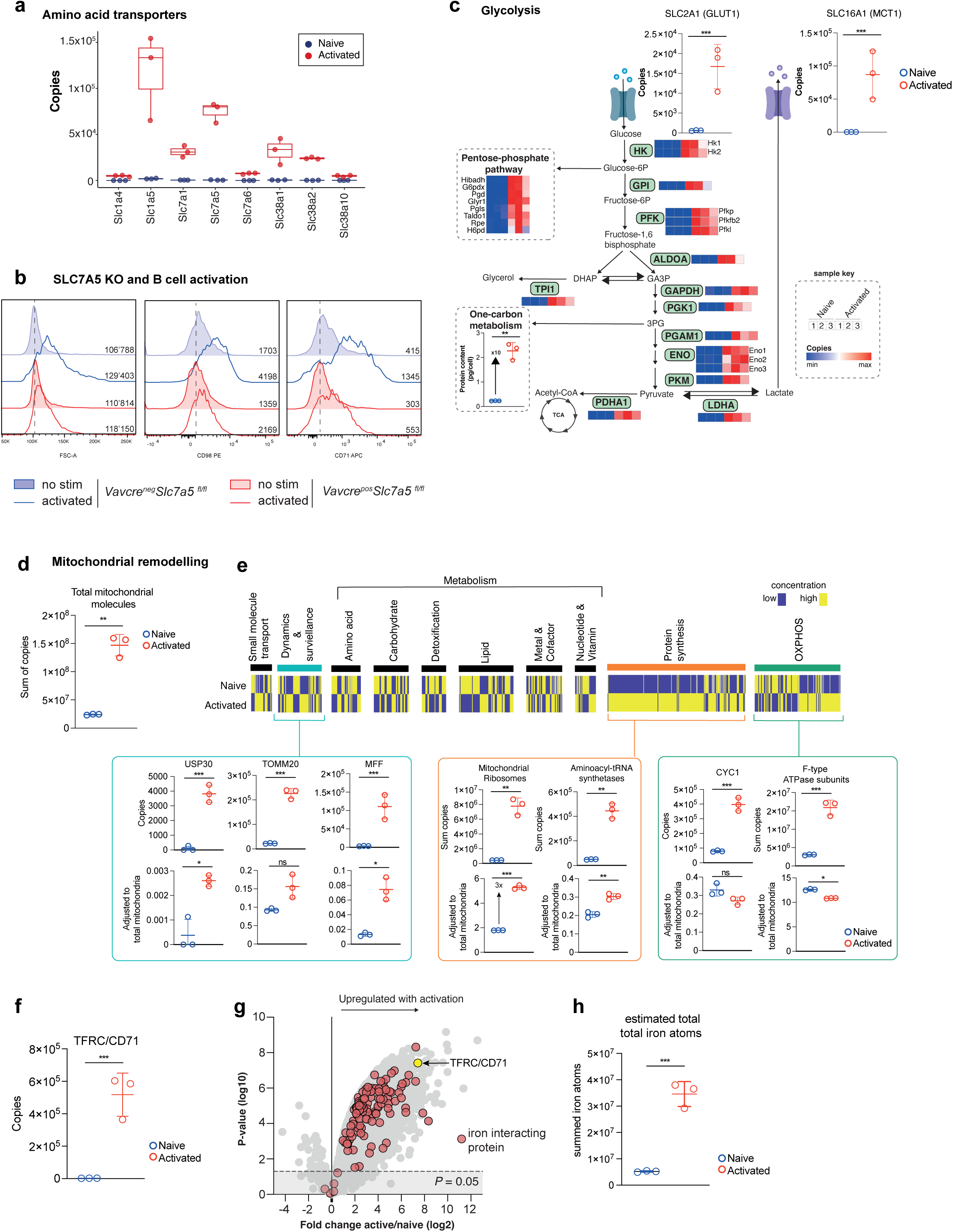
B cell activation triggers metabolic and mitochondrial remodelling. **(a)** Boxplot showing the protein copy numbers per cell of a selection of amino acid transporters. **(b)** The impact of deleting the amino acid transporter SLC7A5 on B cell activation. Using flow cytometry, forward scatter and the expression of CD98/SLC3A2 and CD71 was analysed in Vav cre^+^ Slc7a5^fl/fl^ (SLC7A5 KO; red) and Vav cre^-^ Slc7a5^fl/fl^ (SLC7A5 WT; blue) cells. Splenocytes were activated with anti-IgM and anti-CD40 (activated) or maintained in IL-7 (no stim). **(c)** Schematic representation of the glycolytic pathway. Histograms show the protein copy number per cell for the glucose transporter GLUT1 (SLC2A1) and the lactate transporter MCT1 (SLC16A1). Copy number expression of key enzymes involved in glycolysis and the pentose phosphate shuttle are represented as a heatmap (lower expression is represented by blue and higher expression is represented by red, proteins not detected are shown as grey). Also shown is the total protein content (pg/cell) of proteins involved in one carbon metabolism (GO:0006730). **(d)** >700 proteins were identified as belonging to the mitochondria as per the mouse MitoCarta 3.0 database (Rath et al. 2020b,a) The sum of all mitochondrial protein copy numbers per condition is presented. **(e)** A ratio for the contribution of each mitochondrial protein to the total was calculated by dividing the copy number of each protein by the total mitochondrial molecules. This ratio was used to the plot a heatmap showing relative expression from low (purple) to high (yellow). Heatmap was grouped based on the pathway annotations provided in MitoCarta 3.0. Histograms showing the protein copy numbers per cell, and associated ratio of the total, for proteins in particular pathways are shown. **(f)** Estimated copy numbers per cell for CD71(TFRC) in naïve and activated B cells. **(g)** The expression profile of proteins annotated as iron interacting (red circles) in naive versus activated B cells. **(h)** The estimated total iron atom requirement for those iron interacting proteins identified within naïve and activated B cells. For copy number comparisons of individual proteins statistical significance was derived from two-tailed empirical Bayes moderated t-statistics performed in limma on the total dataset where *** p<0.001. For all other comparisons statistical significance was derived from an unpaired two-tailed t test where * p<0.05, ** p<0.01, *** p<0.001. All data are 3 biologically independent samples except b which is representative of 2 biological repeats. All error bars are mean with s.d.

We next wanted to understand in greater detail the metabolic shift in B cells that occurs upon activation (Boothby & Rickert 2017) . Using estimates of protein copy per cell we mapped glycolytic enzymes and enzymes that drive the pentose-phosphate pathway and one carbon metabolism. Previous studies have questioned the importance of glycolysis in B cell activation, suggesting instead that activated B cells switch on oxidative phosphorylation and the tricarboxylic acid cycle (TCA cycle) but not glycolysis (Waters et al. 2018) . The proteomic data show that expression of the glycolytic machinery, including rate-limiting glucose and lactate transporters (Tanner et al. 2018) as well as glycolytic enzymes are considerably upregulated upon B cell activation (Fig 2c). Indeed, the glucose and lactate transporters SLC2A1 and SLC16A1 were virtually undetectable in naïve cells but were found in tens of thousands of copies upon B cell activation (Fig 2c and Supplementary File 1). The cellular mass of enzymes involved in one-carbon metabolism increased 10-fold in response to B cell activation, suggesting changes in metabolic enzymes that are beyond just scaling with increased cell size (Fig 2c).

Activation induced mitochondrial re-modelling has been well documented in B cells (Boothby & Rickert 2017; Iborra-Pernichi et al. 2024; Waters et al. 2018; Yazicioglu et al. 2023). Using Mitocarta (Rath et al. 2020a) , predicted mitochondrial proteins were annotated within our data set, revealing that total mitochondrial molecules increased approximately 6-fold upon B cell activation (Fig 2d). To explore changes in mitochondrial processes that were independent of activation-induced mitochondrial scaling, we adjusted protein copy numbers by the total mitochondrial protein molecules to determine their relative contribution. This analysis revealed mitochondrial proteins and processes that didn’t just follow the scaled increase in mitochondrial mass (Fig 2e). For example, mitochondrial ribosomes increased 3-fold when adjusted against total mitochondrial molecules, suggesting increased dedication towards mitochondrial protein synthesis in the activating cell. In contrast, molecules involved in oxidative phosphorylation showed a diverse expression pattern with some, including cytochrome C1 and F-type ATPase subunits, remaining at similar concentrations in naïve and activated B cells, suggesting a scaled increase in abundance (Fig 2e).

Having noted significant increases in amino acid, glucose and lactate transporter expression upon B cell activation (Fig 2a and c) we also found considerable upregulation of the transferrin receptor CD71 (TFRC) from around 3000 copies per cell in naïve B cells to approximately 0.5 million copies upon activation (Fig 2f). Using annotations of iron-regulated proteins (Teh et al. 2021) we examined the expression profile of iron-interacting proteins in naïve and activated B cells. We detected 151 iron-regulated proteins in our proteomics data. These included proteins involved in Fe-S cluster synthesis, histone modification (Jmjd family proteins, Kdm family proteins), DNA synthesis (Pol family proteins), and OXPHOS (Fig 2g and Supplementary File 1). To gain a clearer understanding of the cellular requirements for iron, we estimated the total iron-content of the naïve and activated B cells. We assumed that all iron interacting sites were fully occupied and used known values of iron atoms per protein (where possible). Where exact values were not known we used a deliberate under-estimation of 1 iron atom for heme or iron ion interaction, and 2 iron atoms for Fe-S cluster interactions, as described by Teh et al (Teh et al. 2021). Using the protein copy numbers from our data, these calculations showed that the iron demand, or requirement, in naïve B cells is approximately 5 million iron atoms per cell. The demand for iron is increased 7-fold to 35 million iron atoms per cell in activated B cells (Fig 2h). These increased requirements for iron, to drive DNA synthesis, histone modifications and mitochondrial functions, are matched by the increased expression of CD71 (TFRC).

### The selective effects of T cell-dependent versus T cell-independent activation

Next, we compared the impact of T cell-dependent versus T cell-independent activation. Naïve B cells were activated with the TLR4 ligand, lipopolysaccharide (LPS) along with IL4, and the proteome was mapped after 24 hours. The total protein content approximately doubled in response to LPS and IL4 stimulation compared with a 5-fold increase in response to anti-IgM, anti-CD40 and IL4 (Fig 3a). The difference in the scale of proteome remodelling was also reflected in differential expression analysis, with anti-IgM, anti-CD40 and IL4 triggering a median increase in protein abundance greater than 7.4-fold (log2 fold change = 2.9) (Fig 3b) while LPS and IL4 stimulation triggered a median fold change of 2.4 (log2 fold change = 1.3) (Fig 3c). A key question was whether the same proteins and processes were regulated by T cell-dependent versus independent activation. Enrichment analysis of upregulated proteins revealed almost identical biological processes were significantly enriched regardless of the stimulation, with ribosomal and mitochondrial proteins along with DNA replication machinery being consistently increased in abundance (Fig 3d and e). Analysis of the top 500 proteins upregulated in response to either anti-IgM, anti-CD40 and IL4 or LPS and IL4 also revealed that almost all had the same pattern of expression in each stimulation condition (Fig 3f and g). However, important exceptions were identified. Notably, LPS and IL4 triggered the expression of interferon regulated proteins including IFIT1, IFIT2, ISG15 and OAS3 (Fig 3h). Anti-IgM, anti-CD40 and IL4 selectively upregulated a small number of proteins including VAV3, which couples B cell receptor triggering to phosphoinositide 3-kinase signalling and calcium flux(Inabe et al. 2002), tRNA Methyltransferase Activator Subunit 11-2 (TRMT112) which is a key promoter of methyltransferase activity and Transforming Growth Factor Beta 1 (TGFB1) which has pleiotropic effects across immune cells including the regulation B cell activation, proliferation and differentiation. Interestingly, transcription factors that have important roles in regulating B cell fate were also specifically upregulated in response to anti-IgM, anti-CD40 and IL4 but not LPS and IL4. These include aryl hydrocarbon receptor (AHR) and BACH2 which regulate B cell class switching and B cell differentiation(Muto et al. 2010; Vaidyanathan et al. 2017) and EGR2 (Fig 3i). Together these results highlight that the scale of proteome remodelling is drastically different upon T cell-dependent versus T cell-independent activation and key determinants of B cell fate and B cell differentiation are selectively regulated.

**Figure 3.**
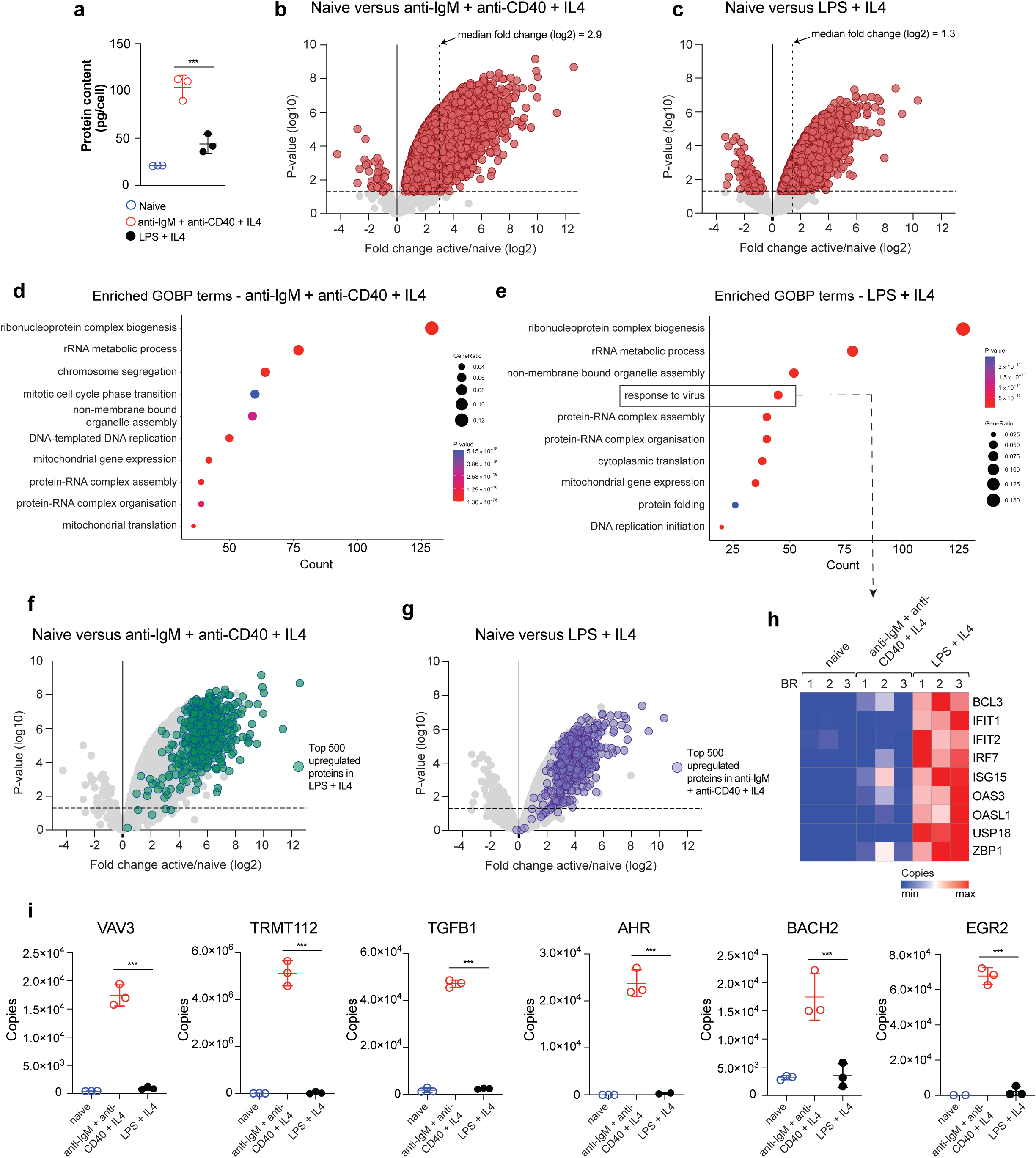
Shared and unique features of T cell-dependent versus T cell-independent B cell activation. **(a)** Total protein content (pg/cell) of naïve B cells and B cells activated for 24 hrs with either anti-IgM + anti-CD40 + IL4 or LPS + IL4. (b and c) Volcano plot showing the differential expression profile of proteins as a consequence of either **(b)** anti-IgM + anti-CD40 + IL4 or **(c)** LPS + IL4 treatment. Data is presented as the log2 fold change of the copy number ratio (activated vs naive B cells) against the inverse significance value (-log10 P value). The median fold change is shown for each data set as a dashed vertical line. (d and e) GO enrichment analysis of proteins significantly upregulated in response to stimulation plus proteins only identified in activated B cells and absent in naive B cells. **(d)** Enriched GO terms in response to anti-IgM + anti-CD40 + IL4 or **(e)** LPS + IL4. Data shown are the top 10 most enriched GOBP pathways. (f and g) Comparative analysis of the expression profile of proteins between two stimulation conditions. **(f)** Top 500 upregulated proteins in response to LPS + IL4 were examined for their expression in response to anti-IgM + anti-CD40 + IL4. **(g)** The top 500 upregulated proteins in response to anti-IgM + anti-CD40 + IL4 were examined for their expression in response to LPS + IL4. **(h)** The expression profile of proteins annotated as ‘response to virus’ across naïve and activated populations. **(i)** The expression profile of a selection of proteins across naïve and activated populations. For b, c, f and g, differential expression analysis and statistical significance was derived from two-tailed empirical Bayes moderated t-statistics performed in limma on the total dataset. For a and i, statistical significance was derived from an unpaired two-tailed t test where *** p<0.001. All data are 3 biologically independent samples. All error bars are mean with s.d.

### The synergistic impact of B cell stimuli

Stimulating B cells with anti-IgM, anti-CD40 and IL4 caused substantial proteome remodelling. To dissect the synergistic effects of these stimuli, B cells were stimulated with anti-IgM, anti-CD40 and IL4 individually and in combinations and their proteomes analysed by single-shot mass spectrometry. Over 7000 proteins were identified across the experiment. As expected, triggering the B cell receptor with anti-IgM caused a down-regulation of the signalling subunits of the B cell receptor, CD79a and CD79b (Supplementary Fig 2a and b) and an upregulation of the nuclear receptor NR4A3 which has previously been shown to be upregulated upon BCR triggering (Supplementary Fig 2c). NFIL3 expression was induced only in conditions in which IL4 was present (Supplementary Fig 2d). Together these results provided confidence in the ability to identify selective effects of stimulation using this data set. Next, we explored the impact of individual and combined stimuli on total protein abundance and the expression profile of proteins driving key cellular processes. B cell stimuli had additive effects on cellular protein abundance, with the addition of IL4 triggering a boost in protein synthesis (Fig 4a). IL4 promotes B cell survival, differentiation and isotype class switching and previous studies have revealed that IL4 induces MYC expression (Klemsz et al. 1989) and drives cholesterol metabolism in B cells (Steach et al. 2024) . Stimulating B cells with IL4 alone caused a modest increase in cell size, MYC expression and protein synthesis (Supplementary Fig 2e-g). However, IL4 in combination with anti-CD40 or anti-IgM induced a significant increase in the abundance of glycolytic enzymes, glucose and lactate transporters and ribosomal proteins (Fig 4b and c). Maximal expression of these molecules was induced in B cells triggered through the BCR, CD40 and in the presence of IL4. Individually, each of these stimuli triggered a downregulation of the cell cycle inhibitor CDKN1B (P27) (Fig 4d). IL4 or anti-CD40 alone had little impact on cell cycle machinery, but together triggered modest increase of cyclins A2, B1 and B2 and CDK2, 4, 6 and 7. Anti-IgM in combination with IL4 or anti-CD40 was clearly the major driver of the expression of the cell cycle machinery (Fig 4d).

**Figure 4.**
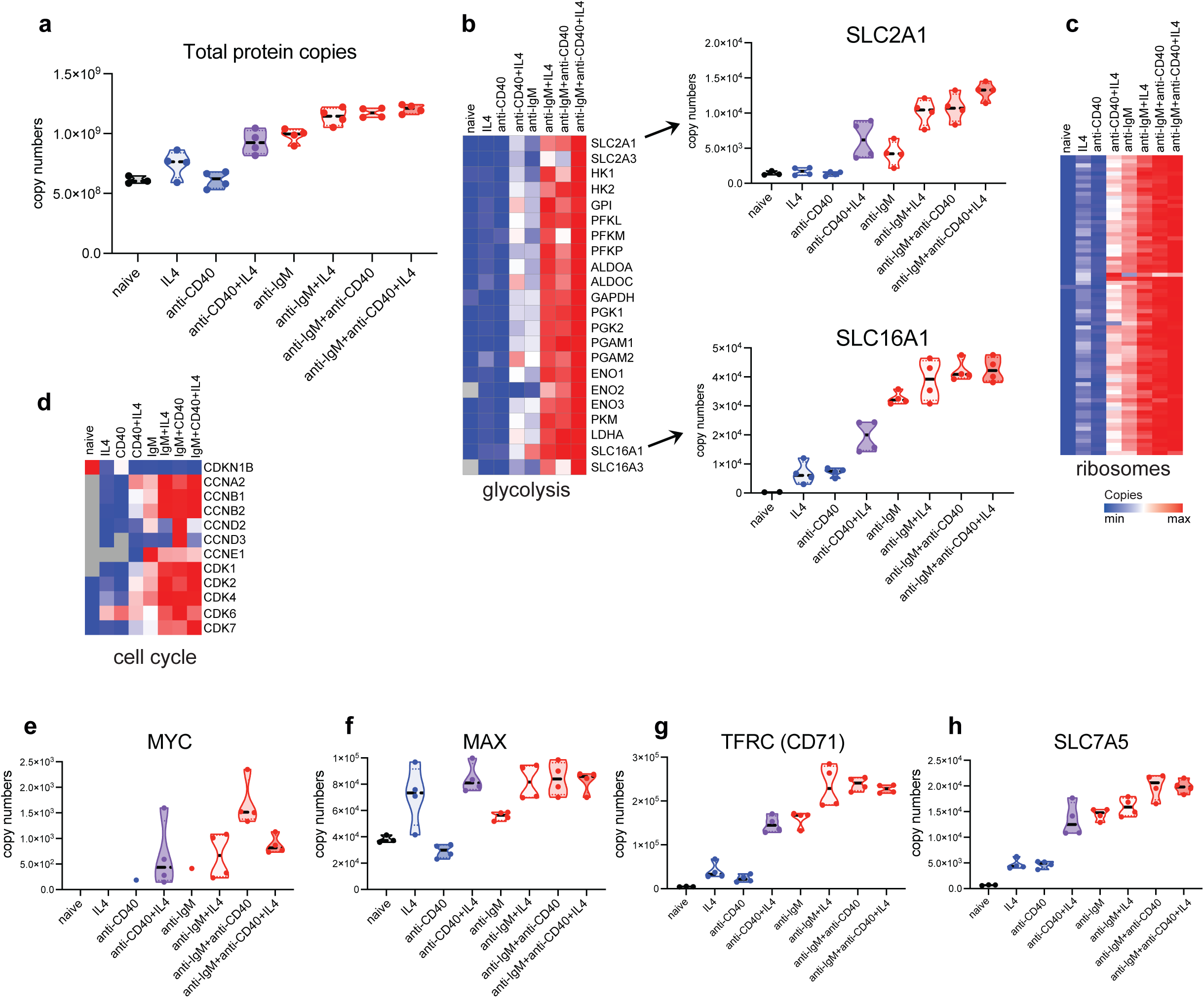
The synergistic impact of B cell stimuli. **(a)** Total protein copies per cell for naïve B cells and B cells activated individually and with combinations of IL4, anti-IgM and anti-CD40. **(b)** Expression profile of proteins belonging to the glycolytic pathway including glucose and lactate transporters. **(c)** The impact of combinations of B cell stimuli on the expression of ribosomal proteins and **(d)** cell cycle machinery. The expression profile of MYC **(e)**, MAX **(f)**, TFRC **(g)** and SLC7A5 **(h)** in response to B cell stimuli. All data are from at least 3 biologically independent samples. All error bars are mean with s.d.

The transcription factor MYC is a central regulator of cellular protein synthesis and growth. As mentioned above, MYC expression was modestly induced by IL4, but its detection was challenging by mass spectrometry, particularly in cells stimulated with either IL4, anti-CD40 or anti-IgM alone. This is likely due to the rapid turnover of MYC within the cell and due to these stimuli triggering small increases in MYC expression. However, combinations of stimuli resulted in increased expression of MYC and its binding partner MAX (Fig 4e and f). B cell receptor and CD40 triggering were required for maximal expression of MYC, supporting previous findings (Luo et al. 2018). A primary function of MYC in lymphocytes is the regulation of amino acid transporter expression(Marchingo et al. 2020) . Known MYC regulated transporters, TFRC (CD71) and the methionine transporter SLC7A5, were maximally expressed in response to co-stimulatory signals (Fig 4g and h).

### mTORC1 activity drives protein biosynthesis and the machinery for fuelling B cells

Together, the B cell activation data reveals the complex interplay between B cell stimuli in driving protein synthesis and the remodelling of the core cellular machinery. Given the central role that mTORC1 plays in regulating diverse cellular responses including lymphocyte growth and differentiation (Chi 2012; Goul et al. 2023; Howden et al. 2019; Iwata et al. 2017; Pollizzi & Powell 2015; Raybuck et al. 2018) we next wanted to explore how mTORC1 regulates the protein biosynthetic capacity of activating B cells and the impact of blocking mTORC1 activity on B cell proteome remodelling. To selectively block mTORC1 during B cell activation, cells were treated with rapamycin and mTORC1 activity monitored through the phosphorylation of ribosomal protein S6, a downstream target of active mTORC1 signalling. Blocking mTORC1 activity with rapamycin during B cell activation reduced S6 phosphorylation to levels similar to those observed in naïve B cells (Fig 5a). Rapamycin treated cells were significantly smaller in size and had approximately half the total cellular protein mass when compared to control activated cells (Fig 5b and 5c). Further, B cells activated with rapamycin exhibited a considerable reduction in rates of protein synthesis compared to control activated cells, as measured by the incorporation of an analogue of puromycin into elongating protein chains in the ribosome (Fig 5d). We sought to understand how the inhibition of mTORC1 was driving the reduction in rates of protein synthesis and consequent decrease in protein mass. Proteomic analysis of cells treated with rapamycin revealed that >4700 proteins had reduced abundance when mTORC1 activity was blocked, while only 28 proteins were found at significantly increased levels (Fig 5e). GO term enrichment analysis revealed that mTORC1 inhibition had a profound effect on DNA replication machinery and mitochondrial translation (Fig 5f). A closer examination revealed that ribosomal protein mass and mitochondrial protein mass were both significantly reduced in response to blocking mTORC1 (Fig 5g). Transcription factors were among some of the most rapamycin-sensitive proteins in B cells. IRF4, a critical regulator of B cell reprogramming during early activation (Patterson et al. 2021) , was found at significantly lower levels in rapamycin treated cells (Fig 5h). IRF8 expression was also significantly impaired when mTORC1 activity was blocked. IRF8 has previously been shown to be important during B cell development and B cell germinal centre responses (Lee et al. 2006; Wang et al. 2008, 2019) , although the role of IRF8 during B cell activation remains unclear. Our proteomic data also revealed high expression of AHR exclusively in activated B cells, which was found at significantly reduced levels, along with the Aryl Hydrocarbon Receptor Nuclear Translocator (ARNT), by rapamycin treatment (Fig 5i). AHR deletion has been shown to impact B cell proliferation (Villa et al. 2017). Interestingly, several cyclin dependent kinases were found at significantly reduced levels in response to mTORC1 inhibition, notably CDK1, which was found at over 120,000 copies per cell in activated B cells, but only 5,000 copies in B cells activated with rapamycin (Supplementary File 1). Together, this data suggests that blocking mTORC1 with rapamycin significantly impacts activation-induced proteome remodelling, reducing the abundance of key proteins involved in ribosomal and mitochondrial mass, transcriptional regulation, proliferation and cell cycle progression.

**Figure 5.**
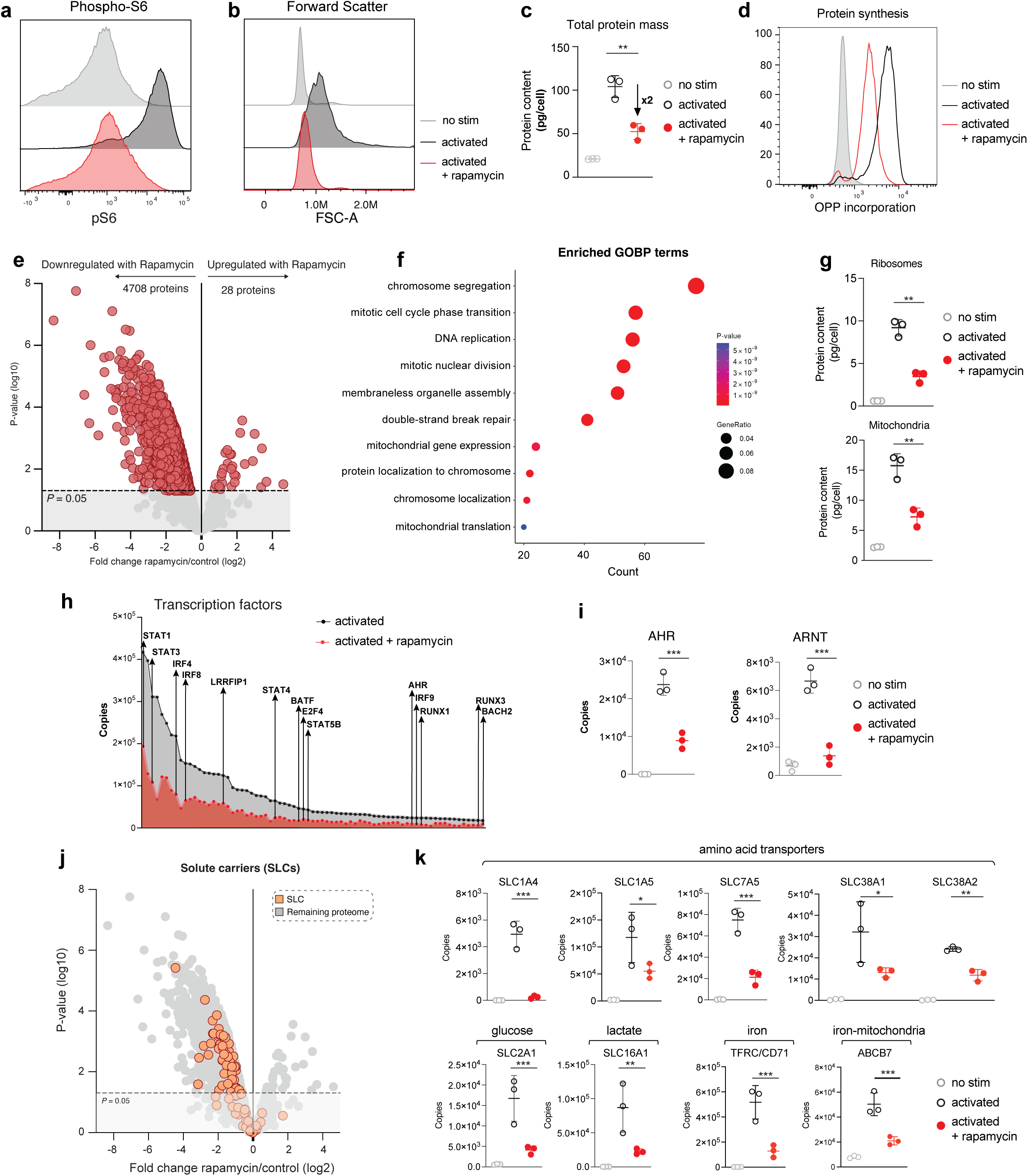
The mTORC1 regulated proteome. **(a)** Phosphorylation of S6 in non-stimulated (grey), activated (black) and activated + rapamycin (red) cells, as determined by flow cytometry. **(b)** Flow cytometry forward scatter (FSC) profile of non-stimulated (grey), activated (black) and activated + rapamycin (red) cells. **(c)** Total protein content (pg/cell) of naïve and 24h activated B cells +/- rapamycin. **(d)** Protein synthesis determined by O-propargyl-puromycin (OPP) incorporation into elongating protein chains and flow cytometry. **(e)** Volcano plot showing the differential expression profile of proteins as a consequence of rapamycin treatment. Data is presented as the log2 fold change of the copy number ratio (rapamycin treated vs untreated activated B cells) against the inverse significance value (-log10 P value). Proteins were considered significantly differentially expressed if the fold change value >1.5-fold in either an up- or down-regulated direction and if the P-value < 0.05 (significance cut off represented as a dashed line). **(f)** GO enrichment analysis of proteins significantly downregulated plus proteins only identified in activated B cells and absent in rapamycin-treated B cells. Data shown are the top 10 most enriched GOBP pathways. **(g)** Total protein content (pg/cell) of proteins belonging to mitochondria (GOCC:0005739) and ribosomes (KEGG 03010). **(h)** Average copy number expression of a subset of transcription factors presented as a line plot between activated B cells +/- rapamycin. Key transcription factors controlling B cell identity and differentiation are shown. **(i)** Protein copy numbers for the Aryl Hydrocarbon receptor (AHR) and the Aryl hydrocarbon nuclear translocator (ARNT) in activated B cells +/- rapamycin. **(j)** Expression profile of solute carriers (SLC) proteins (orange) in response to rapamycin treatment. Log2 fold-change (rapamycin treated vs untreated activated B cells) is plotted against the inverse significance value (-log10 P value). P-value <0.05 represented as a dashed line. **(k)** Protein copy number per cell for a selection of key nutrient transporters for each condition: naïve, 24h activated and rapamycin-treated B cells. For e, i, j and k, differential expression analysis and statistical significance was derived from two-tailed empirical Bayes moderated t-statistics performed in limma on the total dataset where * p<0.05, ** p<0.01, *** p<0.001. For c and g, statistical significance was derived from an unpaired two-tailed t test where ** p<0.01. All data are 3 biologically independent samples. All error bars are mean with s.d. Data presented in d is representative of 3 biological replicates.

mTORC1 inhibition also caused reduced expression of solute carrier proteins (Fig 5j) including the glucose transporter SLC2A1, the lactate transporter SLC16A1 and the System L amino acid transporter SLC7A5 (Fig 5k). The transferrin receptor CD71 and the mitochondrial iron transporter ABCB7 were also significantly reduced in abundance in mTORC1 inhibited cells (Fig 5k). To explore the mTORC1 impact on amino acid transport further, we interrogated the System L amino acid transport capacity of B cells using a sensitive, flow cytometry based assay (Sinclair et al. 2018a). Uptake of the fluorescent SLC7A5 substrate kynurenine was impaired in B cells activated in the presence of rapamycin, orthogonally validating our proteomics data showing reduced levels of SLC7A5 abundance (Fig 6a, 6b, 6c). Flow cytometry analysis also revealed reduced abundance of CD98/SLC3A2 (Fig 6d, 6e) and the transferrin receptor CD71 (Fig 6f, 6g). We next assessed the expression of iron interacting proteins in response to mTORC1 inhibition (Fig 6h) and found that many were considerably reduced in abundance including dioxygenase enzymes ALKBH1-8, a selection of cytochromes, NDUF molecules that are critical for oxidative phosphorylation along with components of the DNA replication fork complex (Fig 6h). Taken together, these data suggest that mTORC1 plays a crucial role in regulating nutrient uptake in B cells, including the regulation of iron transporters and expression of iron interacting molecules.

**Figure 6.**
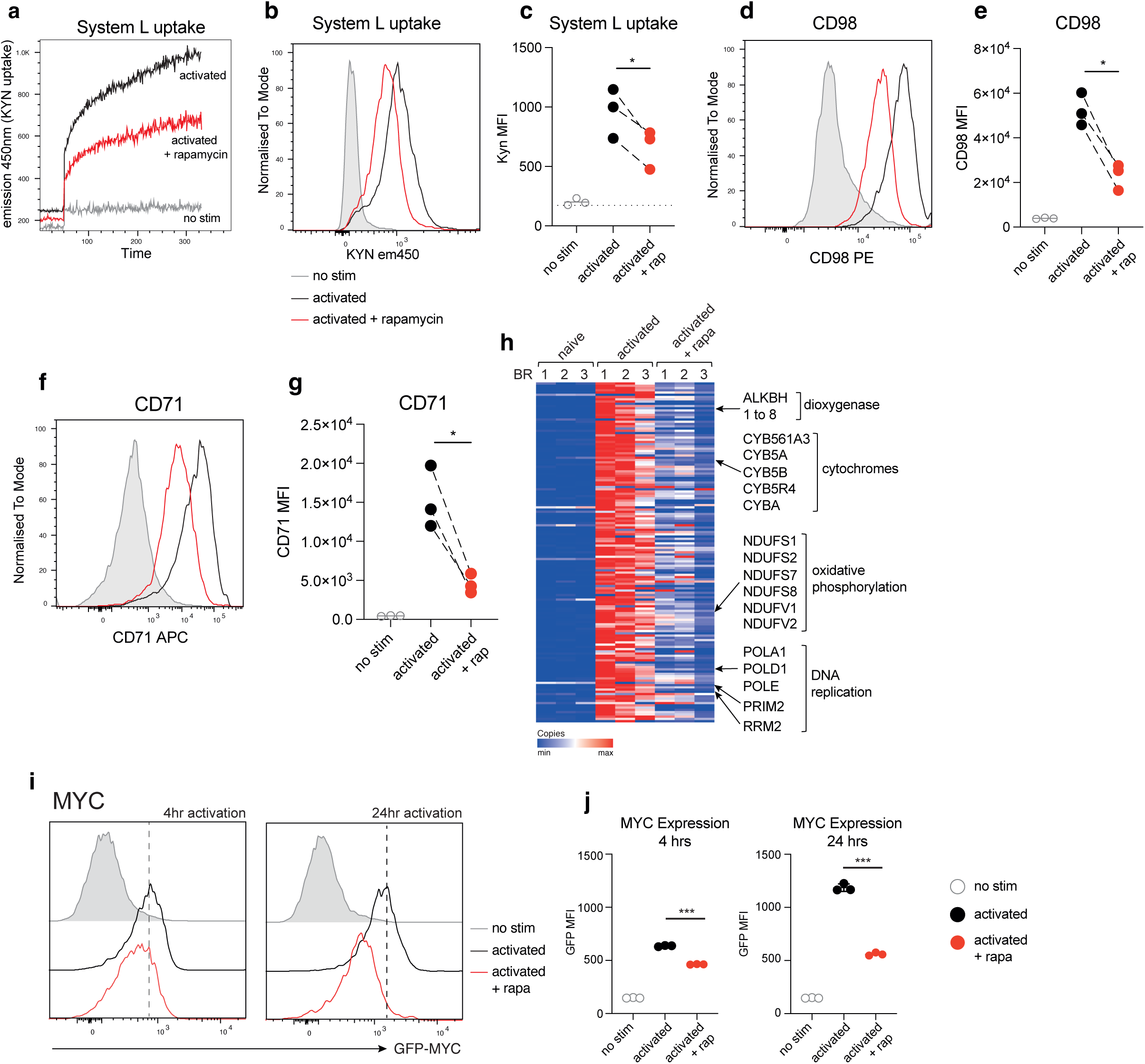
mTORC1 activity regulates amino acid uptake, iron-interacting proteins and MYC expression. **(a)** Live flow cytometric monitoring of System L-dependent uptake in activated B cells +/-rapamycin. B cells were activated for 24 hours before the addition of the System L substrate kynurenine (KYN) and fluorescence emission (450 nm) monitored over time. Unstimulated cells were included as a control. **(b and c)** KYN uptake for 4 minutes and then fixed and analysed by flow cytometry. **(d and e)** The expression of CD98/SLC3A2 in non-stimulated and activated B cells +/- rapamycin. **(f and g)** The expression of CD71 in non-stimulated and activated B cells +/- rapamycin. **(h)** The expression profile of iron-interacting proteins in naive, activated and activated + rapamycin B cells. A selection of iron-interacting protein categories are labelled on the heat map. **(i and j)** The impact of rapamycin treatment on MYC expression in activating B cells. B cells from GFP-tagged MYC mice were activated +/- rapamycin and MYC-GFP expression monitored at 4hrs and 24hrs by flow cytometry. Statistical significance was derived from an unpaired two-tailed t test where * p<0.05, *** p<0.001. For all experiments, 3 biologically independent samples were analysed.

### mTORC1 regulates MYC expression in B cells

We questioned whether mTORC1 driven effects on nutrient transport in B cells could in part be explained by an impact on MYC abundance. MYC is a master regulator of cellular metabolism and growth in T cells(Marchingo et al. 2020). We monitored MYC expression in response to mTORC1 inhibition using a MYC-GFP reporter mouse (Nie et al. 2012) . We found that MYC was upregulated upon B cell activation, but MYC expression was significantly reduced in rapamycin treated cells at 4 hours and 24 hours (Fig 6i and 6j), suggesting the mTORC1 activity is required for maximal MYC expression during B cell activation. Having revealed that mTORC1 activity impacts MYC expression in activating B cells we next sought to understand the consequence of directly blocking MYC activity on B cells. To block MYC at the point of activation, B cells were treated with a small molecule inhibitor of MYC, MYCi361, which inhibits the formation of MYC:MAX heterodimers and blocks the ability of MYC to bind DNA (Han et al. 2019) . Mass spectrometry was used to evaluate the impact of MYC inhibition on the B cell protein landscape while flow cytometry of key surface markers was used for validation. B cells activated in the presence of MYCi361 were reduced in size compared to control cells, according to their forward scatter profile (Fig 7a) and total protein copies per cell was also reduced in response to MYC inhibition (Fig 7b). Over 2500 proteins showed reduced abundance when MYC activity was blocked (Fig 7c). Included in those proteins that were reduced in abundance were ribosomal proteins (Fig 7d). In activated B cells ribosomal proteins totalled over 1.5x10^8^ copies per cell while in MYC inhibitor treated cells this reduced to approximately 0.75x10^8^ copies (Fig 7d). Components of the DNA replication fork machinery were also considerably reduced in expression upon MYC inhibition, in some cases more than 5-fold (Table 1). The impact of MYC inhibition on DNA replication proteins was strikingly similar when compared with rapamycin treated cells (Table 1), highlighting the overlap between mTORC1 and MYC regulated proteins in B cells. Enrichment analysis of those proteins downregulated in response to MYC inhibition showed striking overlap in biological processes regulated by MYC and mTORC1, with DNA replication and mitochondrial translation being consistently impacted (Supplementary Fig 3). Analysis of the top 1000 proteins downregulated in response to either rapamycin or MYCi361 revealed an overlap of 30% of these proteins. However, closer examination revealed that almost all of the 1000 most downregulated proteins had the same pattern of expression in response to rapamycin or MYCi361 (Supplementary Fig 3). To support the MYC inhibitor experiments we explored the impact of MYC deletion on activation-induced proteome remodelling in B cells. The homing receptor CD62L was not impacted by MYCi361 or MYC deletion while the activation marker CD69 was slightly elevated in response to MYCi361 or MYC deletion (Fig 7e-j), in agreement with observations in activating MYC null T cells (Marchingo et al. 2020). Whereas the transferrin receptor CD71 (TFRC), whose expression is dependent on MYC expression in CD8 T cells (Preston et al. 2015) and B cells (O’Donnell et al. 2006), showed reduced abundance in response to MYCi361 and MYC deletion (Fig 7k-m). Additionally, expression of iron interacting proteins were among those reduced upon MYC inhibition (Fig 7n). Interestingly, we noted that nutrient transporters SLC7A5 and SLC2A1, which were both downregulated upon mTORC1 inhibition, were not impacted by MYC inhibition or deletion (Fig 7o-r), indicating expression of these transporters in activated B cells is mTORC1 dependent, but MYC independent.

**Figure 7.**
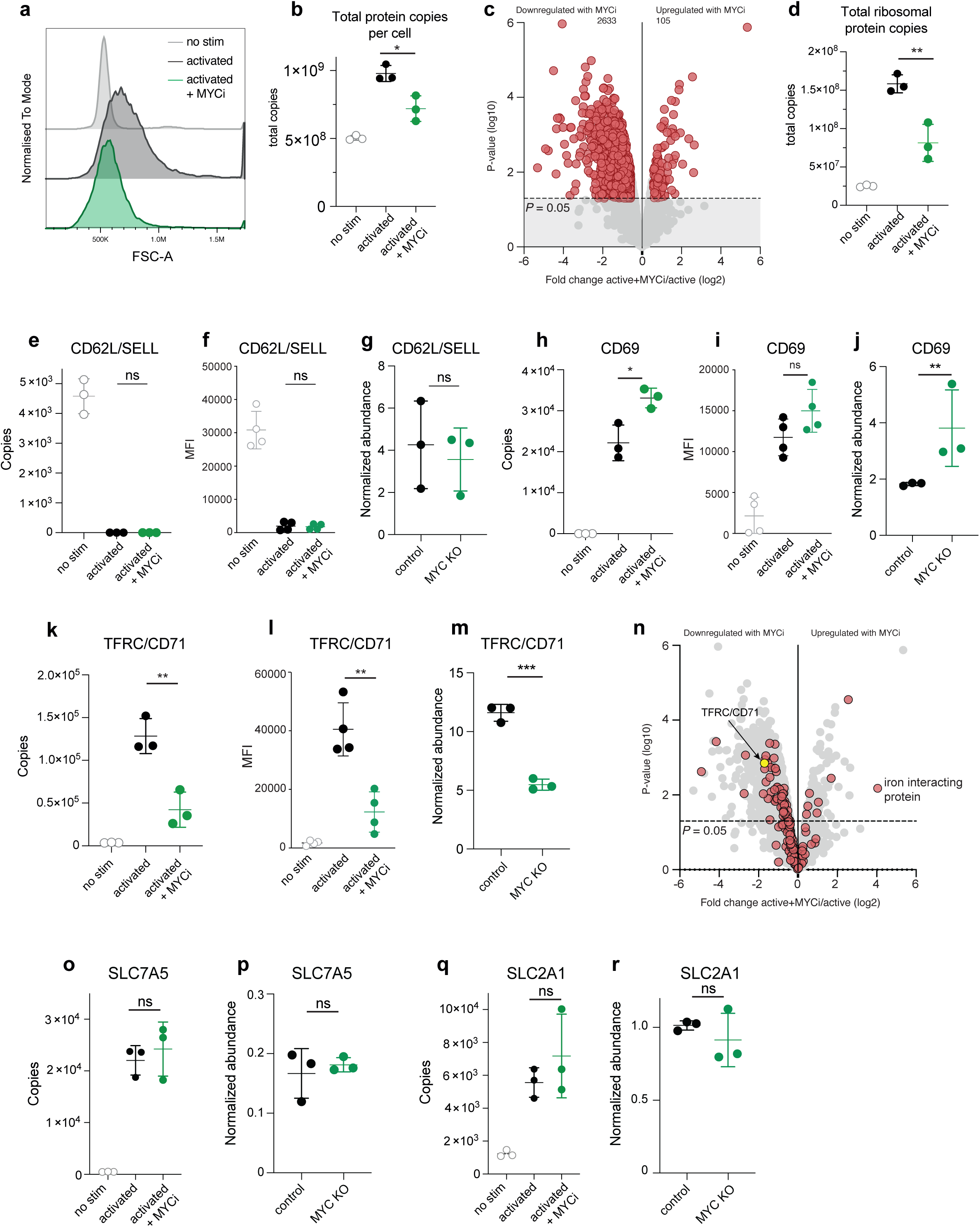
Blocking MYC activity impairs ribosomal protein mass and iron-interacting proteins. **(a)** Flow cytometry forward scatter (FSC) profile of non-stimulated (grey), activated (black) and activated + MYCi361 (green) cells. **(b)** Total protein molecules per cell for naïve and 24h activated B cells +/- MYCi361. **(c)** Volcano plot showing the differential expression profile of proteins as a consequence of MYCi361 treatment. Data is presented as the log2 fold change of the copy number ratio (MYCi361 treated vs untreated activated B cells) against the inverse significance value (-log10 P value). Proteins were considered significantly differentially expressed (highlighted in red) if the fold change value >1.5-fold in either up- or down-regulated direction and if the P-value < 0.05 (significance cut off represented as a dashed line). **(d)** Total ribosomal protein molecules per cell for naïve and 24h activated B cells +/- MYCi361. The impact of MYCi361 or MYC deletion on CD62L (L-selectin), CD69 and CD71 (TFRC) measured by mass spectrometry (**e, g, h, j, k and m**) and flow cytometry (**f**, **i** and **l**). **(n)** Volcano plot showing the expression profile of iron-interacting proteins in response to blocking MYC activity. **(o-r)** The impact of MYCi361 or MYC deletion on the abundance of SLC7A5 and SLC2A1. For b, d, f, i and l statistical significance was derived from an unpaired two-tailed t test where * p<0.05, ** p<0.01. For all other experiments differential expression analysis and statistical significance was derived from two-tailed empirical Bayes moderated t-statistics performed in limma on the total dataset where * p<0.05, ** p<0.01. For all experiments, at least 3 biologically independent samples were analysed. All error bars are mean with s.d.

**Table 1.** The impact of mTORC1 inhibition and MYC inhibition on components of the DNA replication complex. The fold change and p value for a selection of key proteins are provided for activated B cells +/- rapamycin and +/- MYCi361. Statistical significance was determined using a two-tailed empirical Bayes moderated t-test performed in limma on the total dataset.

| Protein | Fold change<br>activ + rapa/activ | p value | Fold change<br>activ + MYCi/activ | p value |
| --- | --- | --- | --- | --- |
| LIG1 | 0.26 | <0.01 | 0.22 | <0.01 |
| MCM4 | 0.25 | <0.001 | 0.23 | <0.001 |
| MCM5 | 0.3 | <0.001 | 0.24 | <0.001 |
| MCM6 | 0.39 | <0.01 | 0.24 | <0.001 |
| MCM7 | 0.27 | <0.001 | 0.23 | <0.001 |
| PCNA | 0.33 | <0.001 | 0.39 | <0.01 |
| POLA1 | 0.17 | <0.01 | 0.16 | <0.001 |
| POLD1 | 0.43 | <0.01 | 0.57 | <0.05 |
| POLE | 0.1 | <0.001 | 0.09 | <0.01 |
| RFC3 | 0.33 | <0.001 | 0.55 | <0.05 |

### Iron availability is essential for B cell growth and protein synthesis

Receptor mediated endocytosis through the transferrin receptor CD71 is the primary method by which lymphocytes take up iron from their extracellular environment. Mutations in CD71 are linked with impaired lymphocyte function and immunodeficiency(Jabara et al. 2016) . Having found that CD71 expression is regulated by MYC and mTORC1 activity and SLC7A5 expression, we next tested whether iron availability regulates activation-induced B cell growth and protein synthesis. Using the iron chelator deferiprone (3-Hydroxy-1,2-dimethyl-4(1H)-pyridone) B cells were activated for 24 hours in iron depleting conditions and monitored by flow cytometry to assess cell size, activation and protein synthesis. Iron chelation impaired the increase in cell size seen upon B cell activation, with increasing concentrations of deferiprone corelating with decreased cell size (Fig 8a-c). However, B cells were still able to upregulate the activation marker CD69 in iron limiting conditions (Fig 8d-e). Protein synthesis was significantly increased in activated B cells when compared with IL4 maintained cells but dropped with increasing concentration of the iron chelator (Fig 8f-g). Higher concentrations of deferiprone (100μM and 300μM) brought levels of protein synthesis down to levels seen in IL4 maintained cells. Together this data shows that iron availability is a critical regulator of activation-induced cell growth in B cells.

**Figure 8.**
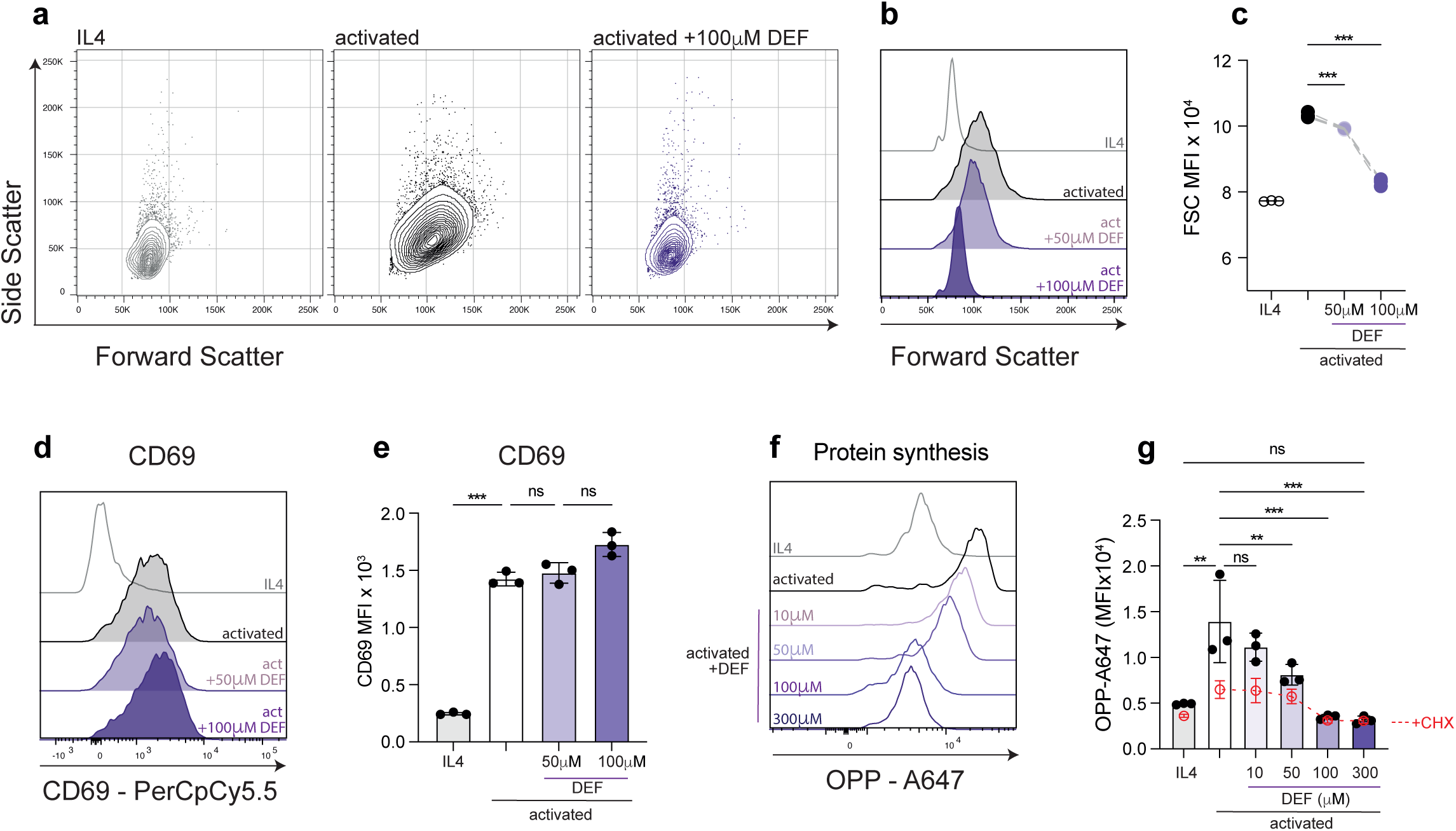
Iron availability is critical for activation-induced B cell growth and protein synthesis. **(a)** Flow cytometry forward and side scatter profile of B cells after 24 hours either maintained in IL4 (1ng/ml) or activated with anti-IgM, anti CD40 and IL4 (10ng/ml) +/- the iron chelator deferiprone. **(b and c)** Forward scatter profile of B cells activated for 24 hours as described above. **(d and e)** CD69 expression in B cells after 24 hours either maintained in IL4 (1ng/ml) or activated with anti-IgM, anti CD40 and IL4 (10ng/ml) +/- deferiprone. **(f and g)** The impact of deferiprone on B cell protein synthesis. B cells were activated for 24 hours and protein synthesis measured by OPP incorporation and labelling with Alexa 647-azide using a copper-catalysed click-chemistry reaction. Cycloheximide (CHX) was included as an inhibitor of cytosolic protein synthesis. For all experiments 3 biologically independent samples were analysed. All error bars are mean with s.d. Data in a, b, d and f are representative of 3 biological replicates. Statistical significance was derived from one way ANOVA where * p<0.05, ** p<0.01, *** p<0.001.

## Discussion

In this work we have mapped the protein landscape of naïve B cells and their response to activation through the B cell receptor, CD40 and the IL4 receptor. Our data reveals the scale of proteome remodelling triggered by B cell activation and the dramatic reconfiguration of cellular compartments including mitochondria and ribosomes. This study aimed to identify the metabolic regulators of B cell activation and protein synthesis. One important biological insight was the identification of the repertoire of amino acid transporters that are expressed upon B cell activation. Deleting one of these transporters, SLC7A5 which transports methionine, tryptophan and leucine, was catastrophic for B cell activation, further highlighting the critical role of this transporter in B cell responses. SLC7A5 has previously been shown to be important during B cell development, with SLC7A5 deletion mice showing reduced B1 B cell numbers (Sun et al. 2023) . SLC7A5 inhibition impacts B cell differentiation and antibody secretion and negatively impacts mTORC1 activity(Torigoe et al. 2019) while deletion of SLC3A2, which forms a heterodimer with transporters including SLC7A5, suppresses B cell proliferation and plasma cell formation(Cantor et al. 2009) . In a complementary study Cheung et al., reveal the importance of SLC7A5 during LPS-mediated B cell activation, with SLC7A5 KO B cells showing impaired activation and proliferation(Cheung et al. 2025) . Together our studies support previous findings showing the fundamental role of a single amino acid transporter in B cell activities.

We also explored the impact of different stimuli on B cell proteome remodelling. The scale of protein changes induced by LPS and IL4 versus anti-IgM, anti-CD40 and IL4, were markedly different. However, the cellular processes and pathways impacted by these two stimulation conditions were largely conserved. Notable exceptions included transcription factors driving B cell fate. An examination of the synergistic impacts of B cell stimuli revealed that glycolytic enzymes, ribosomal proteins and cell cycle machinery were maximally expressed in response to co-stimulation. Interestingly, the expression of nutrient transporters was minimally increased in response to anti-CD40 or IL4 but considerably increased in combination. IL4 is a known driver of protein synthesis in B cells. Our data suggests that this is in part due to the synergistic impact of IL4 inducing the machinery for fuelling B cells along with the machinery for protein synthesis and glycolysis.

To understand the metabolic regulation of B cells we examined the impact of blocking mTORC1 activity during B cell activation. While mTORC1 is firmly established as a key regulator of metabolism and protein synthesis(Ben-Sahra & Manning 2017; Valvezan & Manning 2019) , the mechanisms by which it does this in different cell types and at stages of cellular differentiation, is yet to be fully mapped. In B cells, mTORC1 activity regulates the expression of the machinery for the unfolded protein response prior to B cell differentiation into plasma cells(Gaudette et al. 2020b) and is an important regulator of plasma cell differentiation and humoral immunity(Benhamron et al. 2015; Jones et al. 2016). In the current study we wanted to explore the impact of mTORC1 inhibition during B cell activation. Inhibiting mTORC1 had a considerable impact on protein production during B cell activation, and cells activated in the presence of rapamycin had a reduction in total protein mass of approximately 50% compared to control activated cells. This was also reflected in a significant reduction in ribosomal protein mass in rapamycin treated cells. We found that SLC7A5 expression was regulated by mTORC1, with rapamycin treated cells showing reduced abundance of SLC7A5 and reduced SLC7A5-mediated amino acid uptake. We also found that mTORC1 activity regulates the expression of the transferrin receptor CD71. mTORC1 also regulates the expression of transcription factors that direct B cell activation and differentiation including IRF’s, AHR and MYC. Our findings align with a previous study showing mTORC1 dependent regulation of MYC expression in developing B cells(Zeng et al. 2018) . In T cells, the transcription factor MYC is required for the expression of SLC7A5, along with a selection of other nutrient transporters including CD71(Marchingo et al. 2020). Inhibiting MYC activity in B cells phenocopied many of the mTORC1 inhibition effects including a considerable impact on the cell’s DNA replication machinery, ribosomal protein content and the abundance of CD71. Myc deletion has previously been shown to impact activation induced B cell growth and proliferation(Alboran et al. 2001) and our data suggests this in part may be driven by impacting CD71 expression and the machinery for proliferation and protein synthesis. However, we also revealed unique differences between mTORC1 and MYC inhibition in B cells, with MYC inhibition having little impact on the abundance of SLC7A5 and the glucose transporter SLC2A1. The observation that SLC7A5 expression is MYC dependent in T cells but independent in B cells, suggests cell specific regulation of this key amino acid transporter in lymphocytes. Interestingly, while mTORC1 and MYC are considered master regulators of cellular metabolism, their inhibition did not completely block protein synthesis. Much of the B cell protein landscape is not regulated by either mTORC1 or MYC; B cells still increased their cellular protein content during mTORC1 and MYC inhibition. This highlights that the regulatory pathways for protein synthesis and cell growth are complex, and additional hubs are likely to be important.

Iron has been widely shown to be essential for immune cell function. In B cells, iron availability is required for proliferation and antibody responses(Ned et al. 2003; Stoffel & Drakesmith 2024). Our findings show that mTORC1 activity, SLC7A5 expression and MYC activity all regulate CD71 expression during B cell activation. We questioned whether mTORC1 and MYC mediated proteome remodelling could in large part be attributed to iron availability. Indeed, chelating iron during B cell activation impaired B cell growth and protein synthesis. Our data support iron availability as an important component of the metabolic programme that sustains B cell activation.

In conclusion, this study provides in-depth proteomic maps of naïve and activated B cells and the impact of modulating key signalling pathways during B cell activation. This data provides new insights into the control of protein synthesis and cell growth in B cells and the machinery that drives B cell activation.

### Limitations of the study

This study provides one of the most in-depth analyses of B cell proteome remodelling in response immune stimulation to date. While novel biological insights have been provided including the identification of shared and unique features of mTORC1 and MYC inhibition and the comparison of stimulation conditions, these experiments were performed by culturing primary cells *in vitro*. Future studies could explore the B cell protein landscape in immune challenged mice and dissect B cell subpopulations including those within the germinal center. In addition, this study examined B cell activation in mixed cell cultures from dissociated lymph nodes or spleens. This approach was chosen to limit disturbing cells before stimulation, however, activation and proteome remodelling may be impacted by the activity of other cells within the cell culture. Lastly, this study used rapamycin to block mTORC1 activity. Prolonged exposure to rapamycin may also impact the activity of mTORC2(Sarbassov et al. 2006) , and this should be considered when interpreting results.

## Methods

### Mice

Male wildtype C57BL/6 mice (Charles Rivers) aged between 8 and 12 weeks were used for proteomics experiments examining B cell stimuli and inhibitor treatments and for validation experiments examining protein synthesis, amino acid uptake and marker expression. Myc-eGFP mice(Nie et al. 2012) were used to monitor MYC expression by flow cytometry. VavCre x *Slc7a5^fl/fl^* mice were generated and described previously (Sinclair et al. 2013). All mice were maintained in the Biological Resource Unit at the University of Dundee using procedures approved by the University Ethical Review Committee and under the authorization of the UK Home Office Animals (Scientific Procedures) Act 1986.

For proteomics experiments examining the impact of MYC knockout, Myc-floxed mice (Trumpp et al. 2001) were crossed to Cd19Cre (Rickert et al. 1995) to generate *Cd19Cre Myc^wt/wt^*, *Cd19Cre Myc^fl/wt^ and Cd19Cre Myc^fl/fl^* animals. 10-13 weeks old female mice were used for the experiment. All animal experiments were approved by the Veterinary Office regulations of the Cantone Ticino, Switzerland, and all methods were performed in accordance with the Swiss guidelines and regulations.

### Culturing B cells and cell sorting

For all experiments except MYC KO proteomics, cells were cultured at 37 °C with 5 % CO2 in RPMI 1640 containing glutamine (Invitrogen) and supplemented with 10 % FBS (Gibco), 50 μM β-mercaptoethanol (Sigma) and penicillin/streptomycin (Gibco). For proteomic experiments lymph nodes were extracted from mice, mashed in RPMI media before filtering through a 70 μm cell strainer. For generating a pure population of naïve B cells, lymphocytes were incubated with FC block and then stained with CD19 FITC, CD93 APC and DAPI. CD19+ and CD93- live cells were sorted using a Sony LE-MA900. Sorted cells were washed twice with HBSS and snap frozen in liquid nitrogen and stored at -80 °C until processing for mass spectrometry. For T dependent and T cell independent B cell activation, lymphocytes were suspended in RPMI at a final density of 1.5 million cells/ml and activated for 24 hours in the presence of either 10 μg/ml anti-IgM (AffiniPure F(ab’)₂ fragment goat anti-mouse), 10 μg/ml anti-CD40 (FGK4.5/ FGK45 anti-mouse CD40) and 10 ng/ml IL4 or 20 μg/ml LPS and 10 ng/ml IL4. For experiments exploring the synergistic impact of B cell stimuli, lymphocytes were suspended in RPMI at a final density of 1.5 million cells/ml and activated for 40 hours in the presence of combinations of anti-IgM, anti-CD40 and IL4 at the above concentrations. For rapamycin treatment, cells were activated as described above for 24 hours but in the presence of 20 nM rapamycin for the full duration of activation. For MYC inhibition, cells were activated as above in the presence of 5 μM MYCi361. For all activation experiments subject to proteomic analysis cells were FC blocked and stained with CD19 APC (for activation and rapamycin treated cells) or CD19 APC-Cy7 (for control and MYCi361 treated cells) and DAPI, and live B cells were sorted and collected as described above. Gating strategy used for cell sorting is shown in Supplementary Figure 1. For SLC7A5 KO experiments, splenocytes were activated using 10 μg/ml anti-IgM and 10 μg/ml anti-CD40. For iron chelation experiments lymphocytes were activated for 24 hours as described above but in the presence of deferiprone (3-Hydroxy-1,2-dimethyl-4(1H)-pyridone) at 10 μM, 50 μM, 100 μM and 300 μM. For flow cytometry analysis, samples were analysed on either a Novocyte or LSR Fortessa flow cytometer.

For MYC KO proteomics, splenic follicular (FO) B cells were FACS-sorted (SORP FACSymphony S6, BD Biosciences) for B220^+^/CD19^+^/CD93^-^/CD23^+^/CD2^int^ and seeded at a density of 2x10^6^/ml in complete medium (RPMI, 10% FBS, 1% Penicillin-Streptomycin, 1% MEM NEAA, 1% GlutaMAX, 1% Sodium pyruvate, 0.1% 2-mercaptoethanol, all from Gibco). Cells were activated by anti-IgM (Jackson ImmunoResearch), coated overnight at a concentration of 10 μg/ml, together with recombinant mouse IL4 (PeproTech) added to a concentration of 2 ng/ml and cultured at 37 ⁰C for 24 hours. After stimulation, cells were harvested and washed twice with Hanks’ Balanced Salt Solution (HBSS, Gibco). Cells were then centrifuged and dry pellets were snap frozen in liquid nitrogen and stored at -80 °C prior to lysis for MS.

### Flow cytometry assays for system-L amino acid uptake and protein synthesis

System-L amino acid uptake was measured using a single cell flow cytometry assay described previously (Sinclair et al. 2018b). In brief, after cell surface antibody staining non-stimulated and activated cells +/- rapamycin, were washed in pre-warmed HBSS and resuspended in 200μl HBSS. Cells were treated with either an inhibitor of system L transport 2-amino-2-norbornanecarboxylic (BCH) at a final concentration of 10mM or HBSS as a control. Kynurenine (KYN) was added to the cell suspension at a final concentration of 200μM. For live cell uptakes data was acquired on a flow cytometer immediately following KYN addition.

KYN uptake was also assessed for a fixed time of 4 minutes after which time cells were fixed with paraformaldehyde at a final concentration of 1% and analysed by flow cytometry.

Rates of protein synthesis were measured using a click-chemistry reaction and flow cytometry. B cells were incubated for 20 minutes with O-propargyl-puromycin (OPP, Jena Bioscience), and the incorporation of the aminoacyl-tRNA mimetic into newly synthesized polypeptides was measured by labelling the OPP was with Alexa 647-azide (Invitrogen) using a copper-catalysed click-chemistry reaction (Invitrogen). Cells were analysed using a LSR flow cytometer and analysed with FlowJo software (Treestar).

### Proteomics sample preparation and peptide fractionation

For all proteomics experiments except Myc knockout, cell pellets were lysed in 400 μl of lysis buffer (5% sodium dodecyl sulfate, 50 mM triethylammonium bicarbonate (pH 8.5) and 10 mM tris(2-carboxyethyl)phosphine-hydrochloride). Lysates were shaken at room temperature at 1000 rpm for 5 minutes and then incubated at 95 °C at 500 rpm for 5 minutes. Samples were allowed to cool before being sonicated using a BioRuptor (15 cycles: 30 sec on and 30 sec off) and alkylated with 20 mM iodoacetamide for 1 h at 22 °C in the dark. Protein concentration was determined using the EZQ protein quantitation kit (Invitrogen) and protein cleanup and digestion was performed using S-TRAP mini columns (Protifi). Proteins were digested with trypsin trypsin at 1:20 ratio (enzyme:protein) for 2 hours at 47°C. Digested peptides were eluted from S-TRAP columns using 50mM ammonium bicarbonate, followed by 0.2% aqueous formic acid and 50% aqueous acetonitrile containing 0.2% formic acid. Peptides were dried by speedvac before resuspending in 1% formic acid for fractionation.

For naïve, activated and rapamycin B cell proteomes peptides were fractionated using high pH reverse-phase chromatography as described previously(Damasio et al. 2021). Samples were loaded onto a XbridgeTM BEH130 C18 column with 3.5 μm particles (Waters). Using a Dionex BioRS system, the samples were separated using a 25-min multistep gradient of solvents A (10 mM formate at pH 9 in 2% acetonitrile) and B (10 mM ammonium formate at pH 9 in 80% acetonitrile), at a flow rate of 0.3 ml/min. Peptides were separated into 16 fractions which were subsequently concatenated into 8. The fractions were dried and the peptides were dissolved in 1% formic acid and analyzed by liquid chromatography–mass spectrometry.

For Myc knockout proteomes, cell pellets were resuspended in 4% SDS in 100mM Tris pH 7.6 with 10mM DTT and lysed by sonication in a Bioruptor (Diagenode, 15 cycles, 30s on, 30s off, high mode), followed by a 10 minute incubation at 95°C in agitation. The lysates were alkylated with 50 mM iodoacetamide for 30 minutes at room temperature, cleared by centrifugation for 5 minutes at maximum speed, and proteins were precipitated overnight in 80% cold acetone at -20°C. The next day, proteins were pelleted by centrifugation at 13,000 RPM for 20 minutes at 4°C, after which the pellets were dried and resuspended in 8M urea in 50 mM ammonium bicarbonate by Bioruptor sonication. For each sample, an initial digestion was performed on 100 µg of protein with LysC (Wako Fujifilm, 1:100 w/w) for 2 hours at room temperature, after which the digestion buffer was diluted to 2M urea with 50 mM ammonium bicarbonate and trypsin (Promega, 1:100 w/w) was added for overnight digestion at room temperature. Digestion was stopped in 2% acetonitrile (ACN) and 0.3% trifluoroacetic acid (TFA) and the samples were cleared by centrifugation for 5 minutes at maximum speed. Peptides were purified on C18 StageTips(Rappsilber et al. 2007) , and eluted with 80% ACN, 0.5% acetic acid. Finally, the elution buffer was removed by vacuum centrifugation and purified peptides were resolved in 2% ACN, 0.5% acetic acid, 0.1% TFA for single-shot LC-MS/MS measurements.

### Liquid chromatography mass spectrometry analysis (LC- MS/MS)

For naïve, activated and rapamycin B cell proteomes samples were analysed by data dependent acquisition (DDA). 1.5 μg of peptide was injected onto a nanoscale C18 reverse-phase chromatography system (UltiMate 3000 RSLC nano, Thermo Scientific) and electrosprayed into an Q Exactive™ Plus mass spectrometer (Thermo Fisher) as described previously(Lloyd et al. 2024). The following buffers were used for liquid chromatography: buffer A (0.1% formic acid in Milli-Q water (v/v)) and buffer B (80% acetonitrile and 0.1% formic acid in Milli-Q water (v/v). Samples were loaded at 10 μL/min onto a trap column (100 μm × 2 cm, PepMap nanoViper C18 column, 5 μm, 100 Å, Thermo Scientific) equilibrated in 0.1% trifluoroacetic acid (TFA). The trap column was washed for 3 min at the same flow rate with 0.1% TFA then switched in-line with a Thermo Scientific, resolving C18 column (75 μm × 50 cm, PepMap RSLC C18 column, 2 μm, 100 Å). Peptides were eluted from the column at a constant flow rate of 300 nl/min with a linear gradient from 3% buffer B to 6% buffer B in 5 min, then from 6% buffer B to 35% buffer B in 115 min, and finally to 80% buffer B within 7 min. The column was then washed with 80% buffer B for 4 min and re-equilibrated in 3% buffer B for 15 min. Two blanks were run between each sample to reduce carry-over. The column was kept at a constant temperature of 50°C.

The data was acquired using an easy spray source operated in positive mode with spray voltage at 2.445 kV, and the ion transfer tube temperature at 250°C. The MS was operated in DDA mode. A scan cycle comprised a full MS scan (m/z range from 350-1650), with RF lens at 40%, AGC target set to custom, normalised AGC target at 300, maximum injection time mode set to custom, maximum injection time at 20 ms and source fragmentation disabled. MS survey scan was followed by MS/MS DIA scan events using the following parameters: multiplex ions set to false, collision energy mode set to stepped, collision energy type set to normalized, HCD collision energies set to 25.5, 27 and 30, orbitrap resolution 30000, first mass 200, RF lens 40, AGC target set to custom, normalized AGC target 3000, maximum injection time 55 ms.

To generate proteomes examining the impact of MYC inhibition and the impact of individual and combined stimuli on B cells, peptides were analysed by data independent acquisition (DIA) as described previously (Molina-Gonzalez et al. 2023; Sollberger et al. 2024; Walgrave et al. 2023). 1.5 μg of peptides was injected onto a nanoscale C18 reverse-phase chromatography system (UltiMate 3000 RSLC nano, Thermo Scientific) and electrosprayed into an Orbitrap Exploris 480 Mass Spectrometer (Thermo Fisher). The following liquid chromatography buffers were used: buffer A (0.1% formic acid in Milli-Q water (v/v)) and buffer B (80% acetonitrile and 0.1% formic acid in Milli-Q water (v/v). Samples were loaded at 10 μl/min onto a trap column (100 μm × 2 cm, PepMap nanoViper C18 column, 5 μm, 100 Å, Thermo Scientific) equilibrated in 0.1% trifluoroacetic acid (TFA). The trap column was washed for 3 min at the same flow rate with 0.1% TFA then switched in-line with a Thermo Scientific, resolving C18 column (75 μm × 50 cm, PepMap RSLC C18 column, 2 μm, 100 Å). Peptides were eluted from the column at a constant flow rate of 300 nl/min with a linear gradient from 3% buffer B to 6% buffer B in 5 min, then from 6% buffer B to 35% buffer B in 115 min, and finally to 80% buffer B within 7 min. The column was then washed with 80% buffer B for 4 min and re-equilibrated in 3% buffer B for 15 min. Two blanks were run between each sample to reduce carry-over. The column was kept at a constant temperature of 50 °C.

The data was acquired using an easy spray source operated in positive mode with spray voltage at 2.445 kV, and the ion transfer tube temperature at 250 °C. The MS was operated in DIA mode. A scan cycle comprised a full MS scan (m/z range from 350 to 1650), with RF lens at 40%, AGC target set to custom, normalised AGC target at 300%, maximum injection time mode set to custom, maximum injection time at 20 ms, microscan set to 1 and source fragmentation disabled. MS survey scan was followed by MS/MS DIA scan events using the following parameters: multiplex ions set to false, collision energy mode set to stepped, collision energy type set to normalized, HCD collision energies set to 25.5, 27 and 30%, orbitrap resolution 30,000, first mass 200, RF lens 40%, AGC target set to custom, normalized AGC target 3000%, microscan set to 1 and maximum injection time 55 ms. Data for both MS scan and MS/MS DIA scan events were acquired in profile mode.

For Myc knockout proteomes, peptides were separated on a nanoElute2 HPLC system (Bruker) coupled via a nanoelectrospray source (Captive spray source, Bruker) to a timsTOF HT mass spectrometer (Bruker). 500 ng of peptides per sample were loaded in water with 0.1% formic acid into a column (25 cm long, 75 µm inner diameter, kept at 50°C in a column oven) in-house packed with ReproSil-Pur C18-AQ 1.9 µm resin (Dr. Maisch HPLC GmbH), and eluted over a 60-min linear 2-35% gradient of ACN with 0.1% formic acid at a 300 nl/min flow rate. The mass spectrometer was operated in a data-independent (DIA)-PASEF mode with 100 ms of accumulation and ramp time, covering with 21 mass steps, 25 Da wide, and 1 mobility window, a mass range from 475 to 1000 Da and a mobility range from 0.85 to 1.27 Vs cm-2, with an estimated cycle time of 0.95 s.

### Proteomics data processing and analysis

Data dependent mass spectrometry raw files were searched using MaxQuant(Cox & Mann 2008) (version 1.6.10.43). Data was searched against a hybrid database from Uniprot release 06/2020. This hybrid database comprised of all manually annotated mouse SwissProt entries combined with mouse TrEMBL entries with protein level evidence available and a manually annotated homologue within the human SwissProt database. The false discovery rate was set to 1% for positive identification at the protein and peptide level. Protein N-terminal acetylation, methionine oxidation and deamidation (NQ) were set as variable modifications and carbamidomethylation of cysteine residues was selected as a fixed modification. Match between runs was disabled. Proteins categorized as ‘contaminants’, ‘reverse’ and ‘only identified by site’ were removed. Data independent mass spectrometry raw files were searched using Spectronaut (Biognosys) version 19. Raw mass spec files were searched against a mouse data base (Swissprot Trembl November 2023) with the following parameters: directDIA, false discovery rate set to 1%, protein N-terminal acetylation and methionine oxidation were set as variable modifications and carbamidomethylation of cysteine residues was selected as a fixed modification. Copy numbers were calculated using the proteomic ruler(Wiśniewski et al. 2014) using Perseus software(Tyanova et al. 2016). GO term enrichment analysis was performed using the ORA algorithm from the R Bioconductor package clusterProfiler. The annotation database was set to GO biological process (GOBP) and the organism database used was org.Mm.eg.db. A secondary simplification step was performed to remove redundancies from GO enrichment results (default settings were used with a similarity cut-off of 0.6).

### Statistics and calculations

For proteomics data, protein differential expression analysis was performed using R (v. 4.0.3) and p-values and fold changes were calculated using the Bioconductor package Limma (v 3.46.0)(Ritchie et al. 2015) .

## Resource availability

All proteomics data is provided in Supplementary File 1. Raw mass spec data files and analysis files are available from the ProteomeXchange data repository (http://proteomecentral.proteomexchange.org/cgi/GetDataset) and can be accessed with the following identifier’s: PXD056624 for naïve and activated B cell data +/- rapamycin; PXD057711 for activated B cell data +/- MYCi361; PXD070254 for individual and combined B cell stimuli and PXD074535 for Myc knockout data. Flow cytometry data are available from the corresponding author upon request.

## Acknowledgments

The authors would like to thank the University of Dundee flow cytometry facility for cell sorting support and the University of Dundee Biological Resource Unit. We sincerely thank K. Rajewsky, Mouse Genetics Cologne (MGC) Foundation, and A. Trumpp, German Cancer Research Center, Heidelberg, for sharing mouse models for MYC knockout experiments. We also thank Doreen Cantrell, Dana Cheung and Simon Arthur for helpful discussions. Animations were drawn using BioRender. This research was supported by Wellcome Trust grant 205023/Z/16/Z, and a Wellcome Trust Equipment Award, 202950/Z/16/Z.

## Author Contributions

AJMH, NL, HK, MP and LVS performed experiments. AJMH, LVS, HK, GG and FS designed experiments. OJ, LVS, NL, AACB and AJMH analysed data. LVS, OJ and AJMH wrote the manuscript. All authors read and commented on the manuscript. AJMH conceived the project.

## Supplementary Figures

**Supplementary Figure 1.**
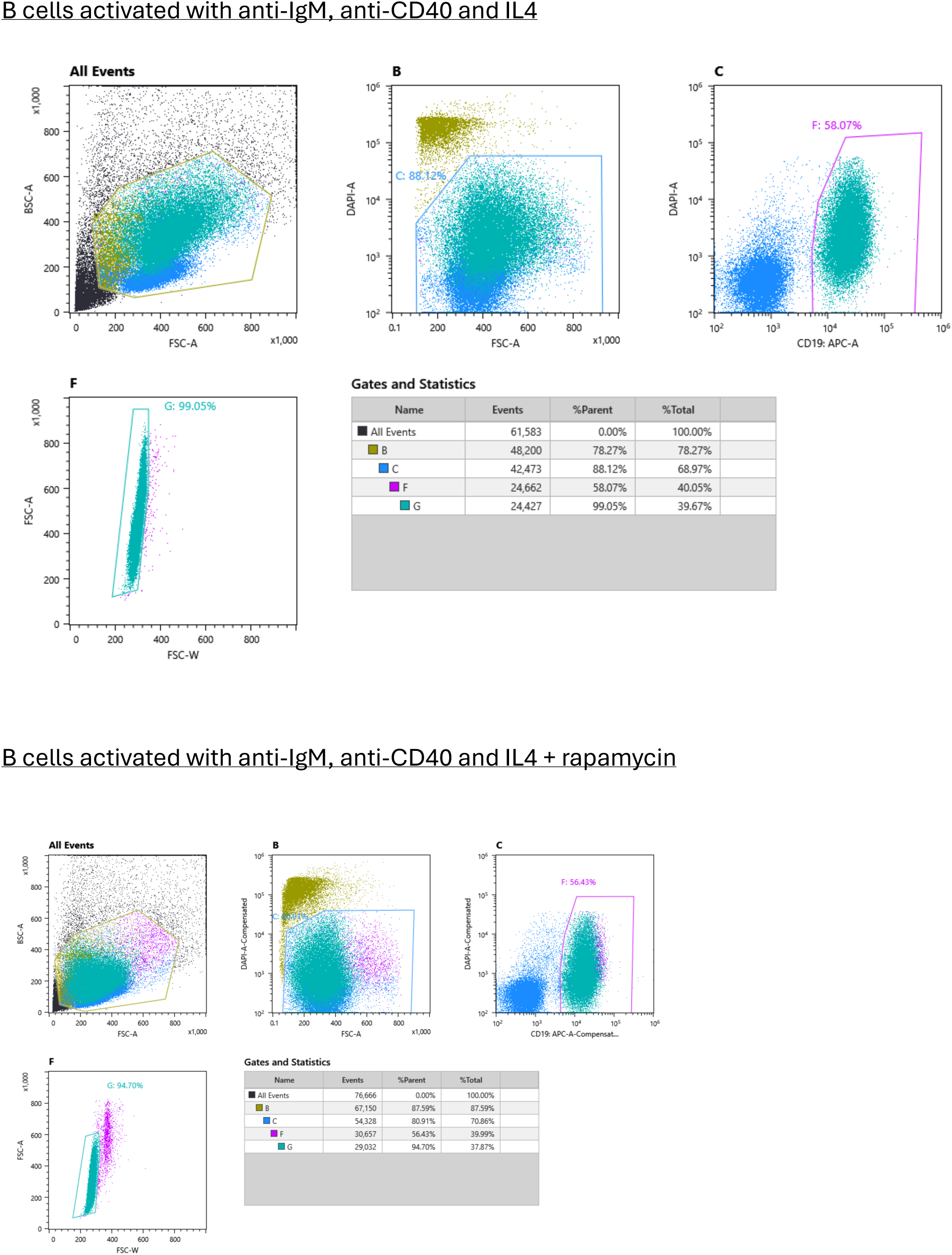
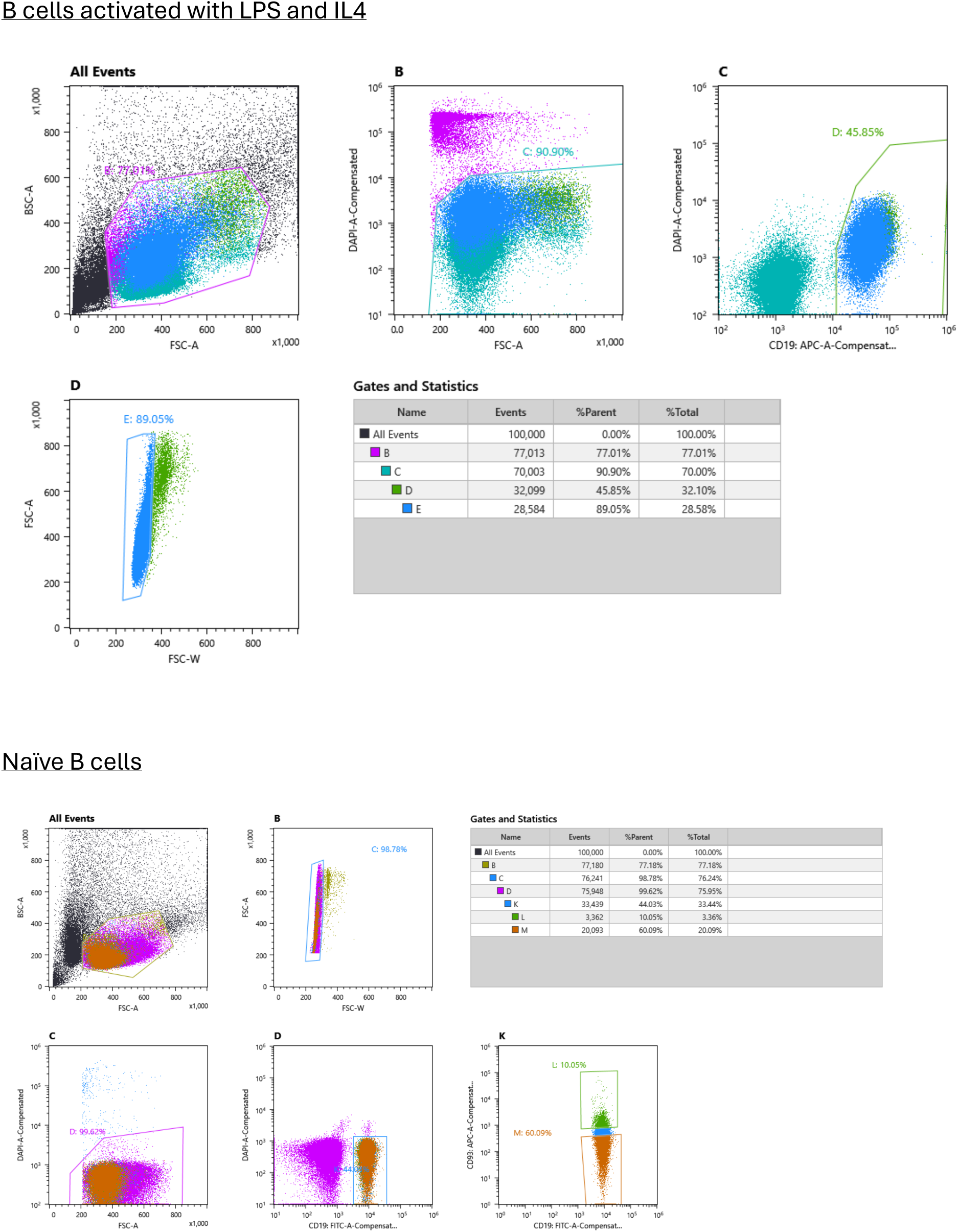
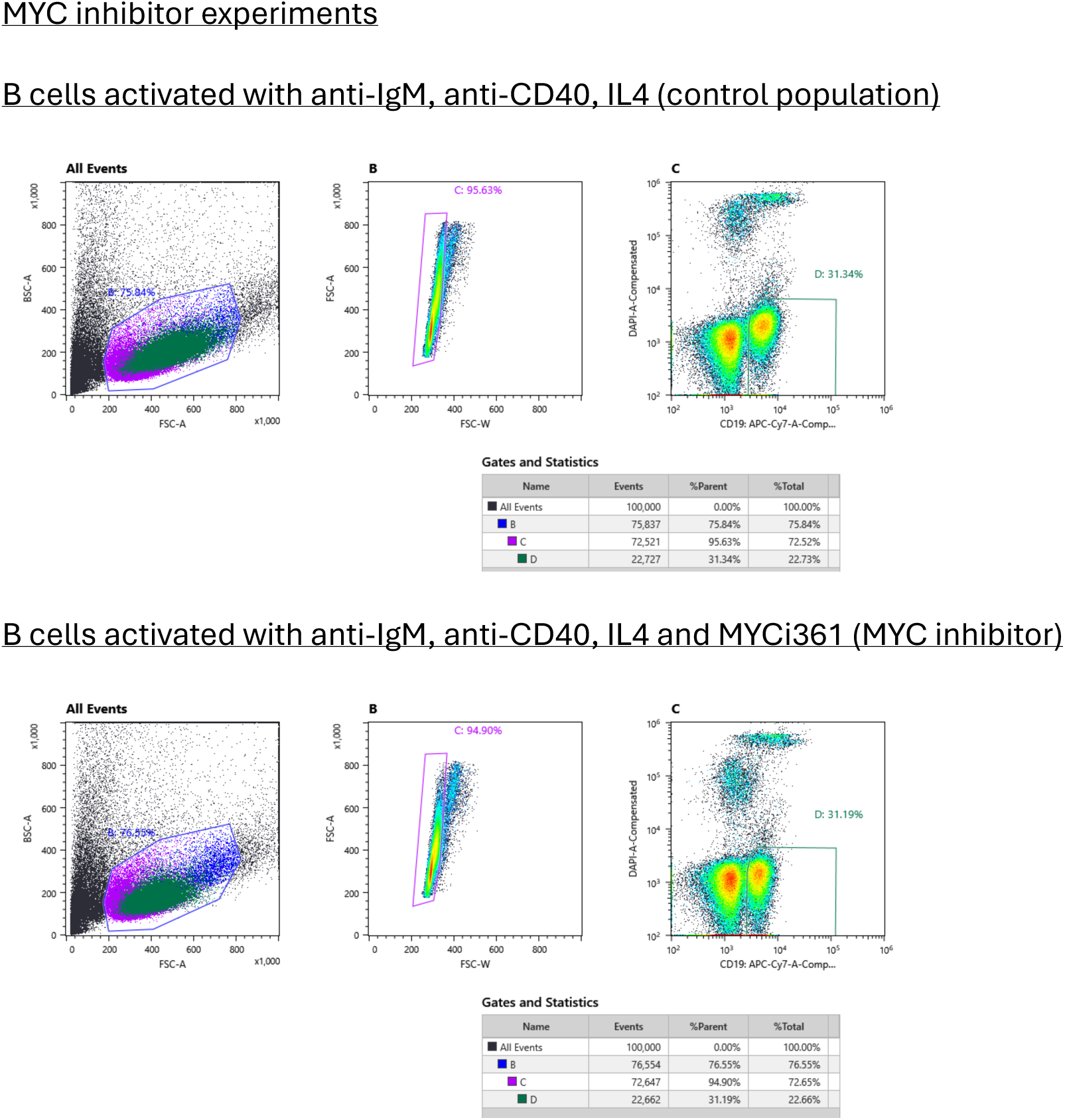
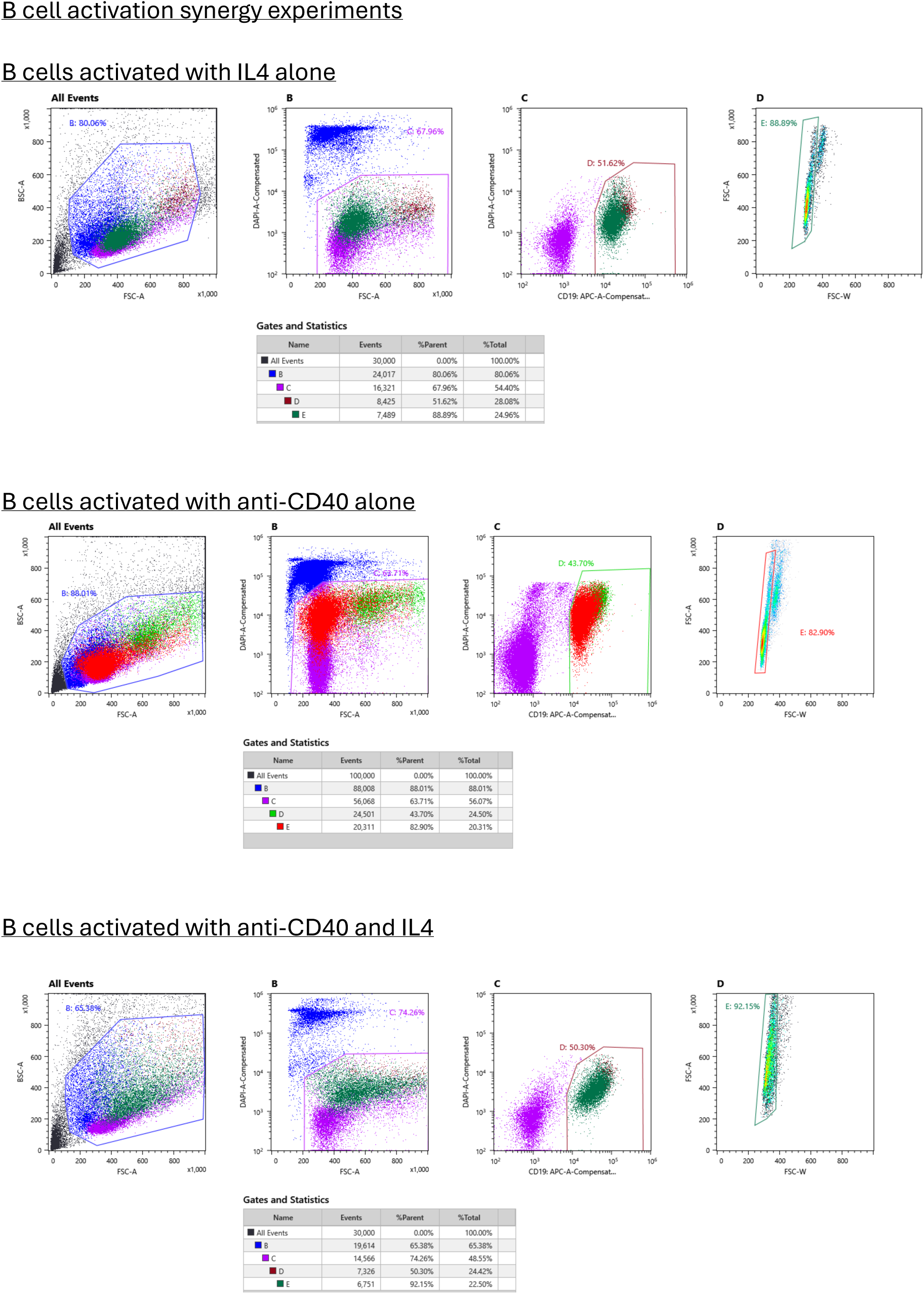
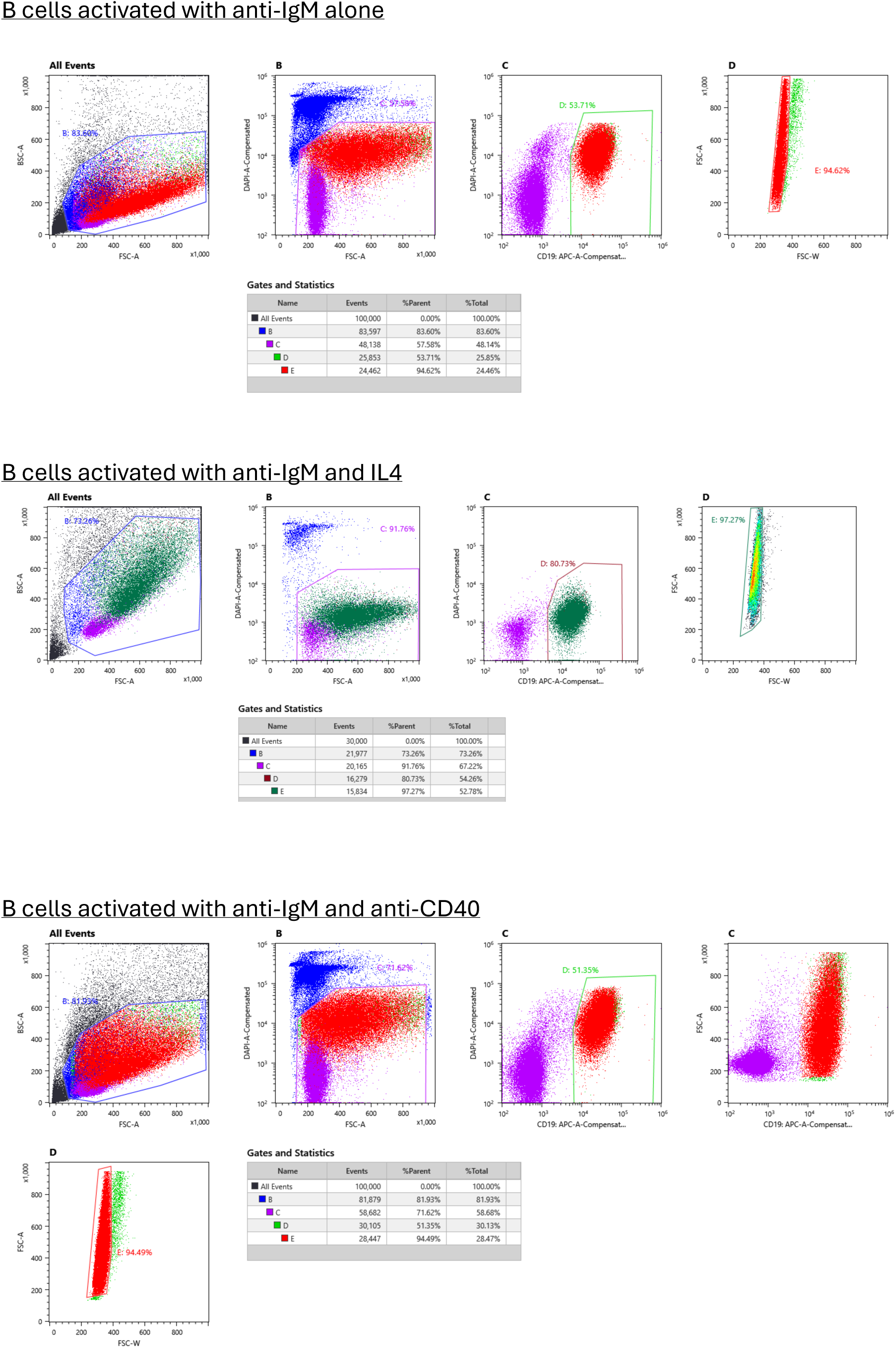
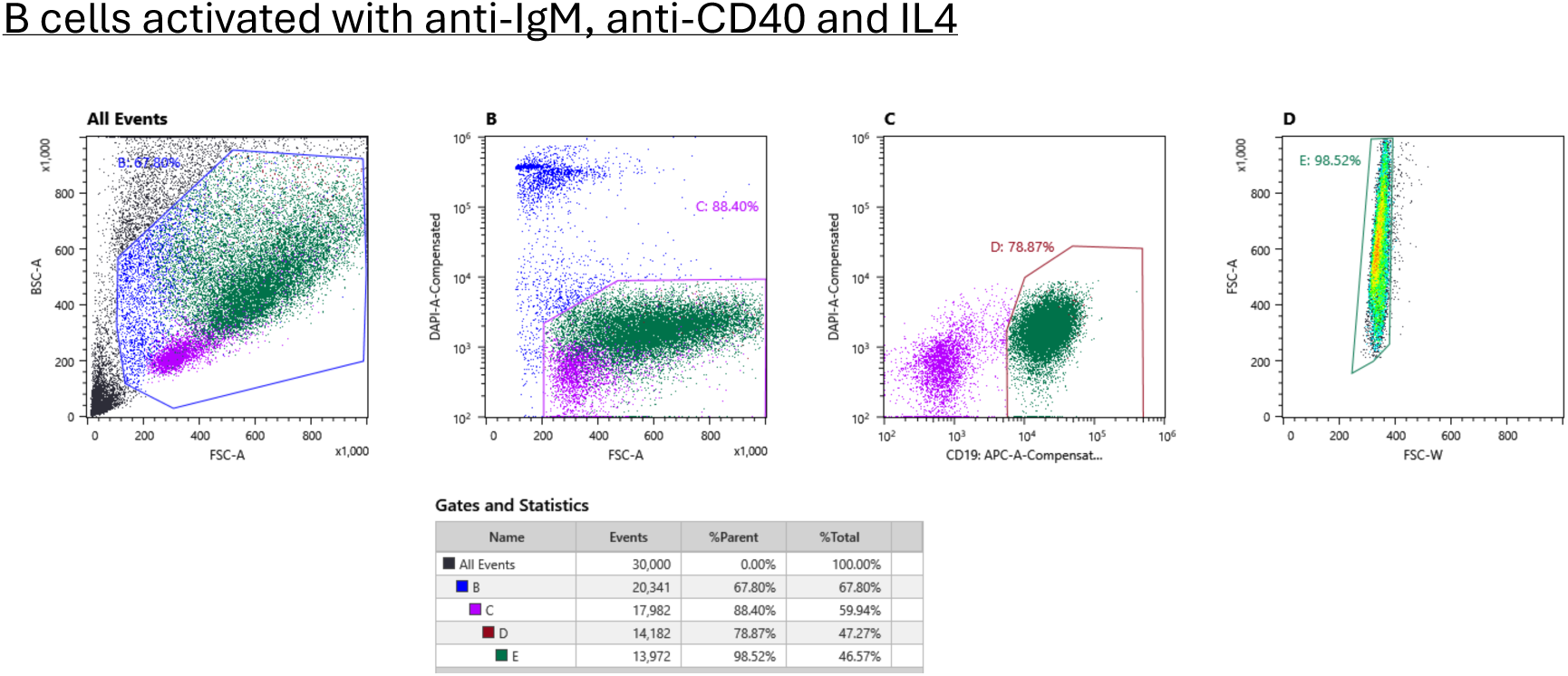
Example gating strategy for cell sorting B cell populations.

**Supplementary Figure 2.**
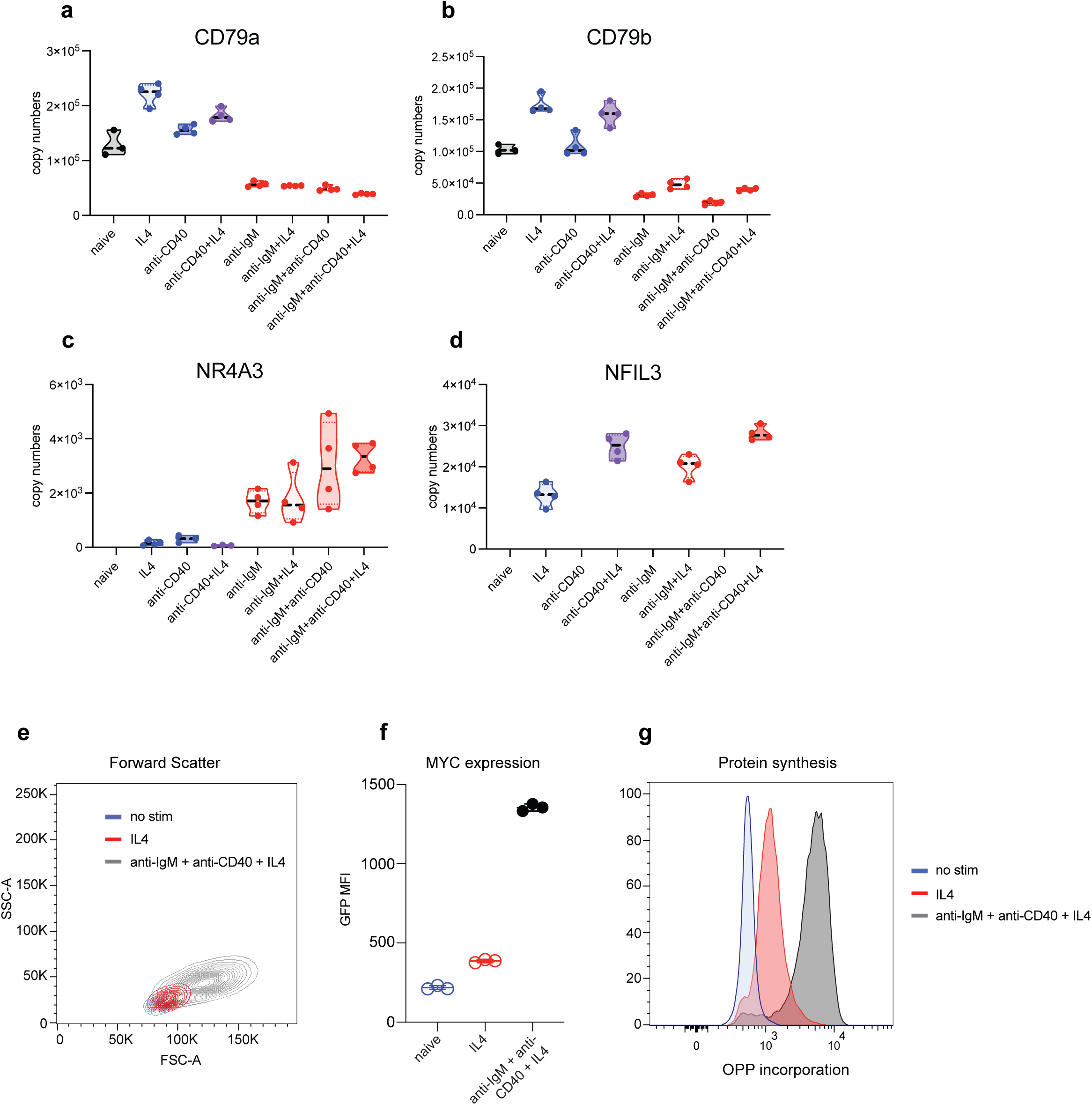
The synergistic impact of B cell stimuli.

**Supplementary Figure 3.**
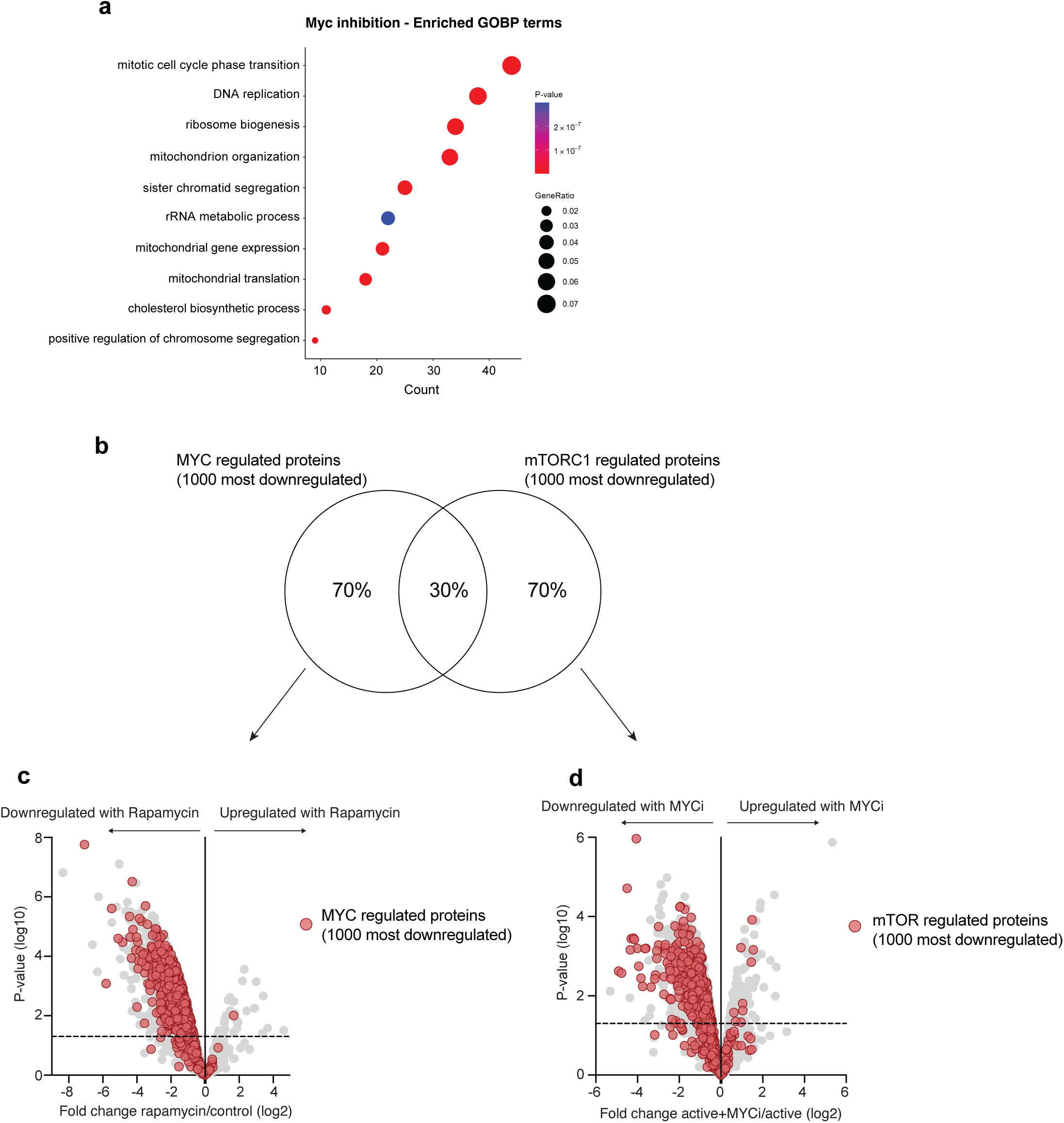
The overlap of mTORC1 and MYC inhibition on activating B cells. **(a)** GO enrichment analysis of proteins significantly downregulated in response to MYC inhibition. Data shown are the top 10 most enriched GOBP pathways. (b) Overlap in protein identifications for the most downregulated proteins in response to MYC inhibition or mTORC1 inhibition. (c and d) Overlap in the expression profile of the most downregulated proteins in response to either MYC or mTORC1 inhibition. Statistical significance was derived from two-tailed empirical Bayes moderated t-statistics performed in limma.

